# Synapse-specific trapping of Syntaxin1a into nanoclusters by the general anesthetic isoflurane

**DOI:** 10.1101/2023.02.27.530184

**Authors:** Adam D. Hines, Victor Anggono, Adekunle T. Bademosi, Bruno van Swinderen

## Abstract

General anesthetics disrupt brain network dynamics through multiple pathways, predominately through post-synaptic potentiation of GABA_A_R and pre-synaptic inhibition of neuroexocytosis. Common clinical general anesthetic drugs, such as propofol and isoflurane, have been shown to interact and interfere with a core component of the exocytic release machinery, Syntaxin1A, to cause impaired neurotransmitter release. Recent in vitro studies however suggest that these drugs to not affect all synapse subtypes equally. We investigated the role of Syntaxin1A in multiple neurotransmitter systems under isoflurane general anesthesia in the adult *Drosophila* brain using live-cell super resolution microscopy and optogenetic readouts of exocytosis. We found that effects of isoflurane anesthesia were neuron specific and only impaired Syntaxin1A activity in excitatory neurons at cholinergic synapses, but not inhibitory GABAergic or glutamatergic neurons. These results indicate that general anesthetics could work by producing successive bouts of inhibition across the brain, by reducing neuroexocytosis from excitatory neurons specifically as well as silencing arousal systems through GABA_A_R potentiation.

## Introduction

General anesthesia is a reversibly induced state of unconsciousness and unresponsiveness that has been used clinically for over 150 years. It has been well understood for over four decades that these drugs interact with proteins in the central nervous system (CNS) to disrupt neuronal activity and silence wake-promoting circuits (Franks, 2008). The best-characterized interaction involves the potentiation of postsynaptic chloride permeable γ-aminobutyric acid type A (GABA_A_) receptors, which act to hyperpolarize wake-promoting neurons and thus promote a loss of consciousness (Franks and Lieb, 1984; Jones et al., 1995; Kim et al., 2020). However, such postsynaptic effects cannot fully account for the complexity of general anesthesia, particularly for the lack of responsiveness during surgery. Accordingly, other protein targets of general anesthetics have been discovered, especially at the pre-synapse, including ion channels (Bertaccini et al., 2014; Koyanagi et al., 2019; Ton et al., 2017; Torturo et al., 2019; Zhou et al., 2019), kinesins (Bensel et al., 2017; Woll et al., 2018), mitochondria (Morgan et al., 2002; Zimin et al., 2018), and the synaptic release machinery (Bademosi et al., 2018b; Herring et al., 2011; Herring et al., 2009; Hines and van Swinderen, 2021; Karunanithi et al., 2020; Troup et al., 2019; van Swinderen et al., 1999; Xie et al., 2013; Zalucki et al., 2015). Although it is known that general anesthetics impair neurotransmitter release and neuronal communication (Bademosi *et al*., 2018b; Herring *et al*., 2011; Herring *et al*., 2009; Karunanithi *et al*., 2020; Speigel and Hemmings, 2021; Wang et al., 2020), the relative contribution of these different pre-synaptic target remains unclear.

The soluble N-ethylmaleimide-sensitive factor attachment protein receptor (SNARE) complex, which comprises syntaxin1A (Sx1a), SNAP25, and VAMP2, is essential for neurotransmitter release (Südhof, 2012). These proteins form a tetramer of tight alpha helical structure that facilitates the docking and fusion of synaptic vesicles with the plasma membrane. Sx1a has been shown in multiple studies to be a target of different general anesthetics, both volatile and intravenous (Bademosi *et al*., 2018b; Herring *et al*., 2011; Herring *et al*., 2009; Hines and van Swinderen, 2021; Karunanithi *et al*., 2020; van Swinderen *et al*., 1999). In the presence of general anesthetic, Sx1a molecules are trapped into non-functional nanoclusters on the presynaptic membrane, potentially preventing them from forming competent SNARE complexes (Bademosi *et al*., 2018b; Hines and van Swinderen, 2021). However, it remains unclear if this effect on Sx1a is observed unilaterally across the CNS, as there is evidence from cell culture experiments that general anesthetics may differentially target excitatory and inhibitory neurotransmitter systems (Baumgart et al., 2015; Speigel and Hemmings, 2021). Furthermore, it is unclear whether general anesthetics also trap other release machinery proteins into non-functional nanoclusters at the presynapse.

In this study, we sought to determine the effects of a volatile anesthetic (isoflurane) on neuroexocytosis and Sx1a function in different neurotransmitter systems (cholinergic, GABAergic, and glutamatergic) in intact brain tissue. Using an *ex vivo* brain preparation of the *Drosophila* fruit fly as a model (Hines and van Swinderen, 2021), we combined optogenetically-induced neuroexocytosis assays (Miesenböck et al., 1998) and single molecule imaging (Zhang et al., 2012) to study how the diffusion and clustering of Sx1a correlates with decreased vesicle fusion dynamics at different synapses. We found that in conjunction with impaired synaptic release, Sx1a molecules become more immobilized in both synaptic and extrasynaptic compartments of cholinergic neurons under isoflurane anesthesia, but only Sx1a proteins at active zones were entrapped into clusters. Notably, we saw no change in exocytosis or Sx1a diffusion dynamics under isoflurane general anesthesia in two inhibitory neurotransmitter circuits (GABA and glutamate), which do not express the key SNARE protein, SNAP25. Together, this provides a model for how presynaptic inhibition of excitatory neurotransmission might contribute to the behavioral effect of general anesthesia.

## Results

### Isoflurane impairs synaptic vesicle exocytosis in the fly brain

To establish the effects of general anesthesia in the *Drosophila* brain, we administered the volatile agent isoflurane to adult flies to induce unconsciousness and a loss of behavioural responsiveness. Flies were exposed to either a 1.5 vol% dose of isoflurane (Zalucki *et al*., 2015) or air as a control for 15 min, before brain dissection and imaging (Figure 1A). To quantify synaptic vesicle exocytosis, we used synaptopHlourin, a pH-sensitive reporter of vesicle fusion to the plasma membrane (Miesenböck *et al*., 1998) (Figure 1B). We expressed the synaptopHluorin in Mushroom Body Output Neurons (MBONs) (Aso et al., 2014), by selecting MBON split-Gal4s to drive the transgene expression in specific MBON subsets that have been mapped to different neurotransmitter systems (cholinergic, GABAergic, glutamatergic, etc.) by the *Drosophila* hemibrain connectome (Scheffer et al., 2020). We started with a set of cholinergic MBONs by using the MB543B-Gal4 driver (Figure 1C,D), as the major excitatory neurotransmitter system in the fly brain is cholinergic (Gu and O’Dowd, 2006). To stimulate synaptic release, we co-expressed the Ca^2+^ permeable channelrhodopsin CsChrimson tagged with mCherry and activated with red light exposure (Figure 1B). SynaptopHluorin was strongly expressed in synapses but not extrasynaptic areas or cell bodies, whereas CsChrimson expression was evident throughout the entire neuron (Figure 1D). All neurons were subjected to a standardised imaging protocol of 10 s baseline, 2 min of 10 Hz CsChrimson activation, and 5 min of recovery post-stimulation (Figure 1A, timeline; Figure S1A; Supplemental Movie 1). We uncovered dynamic synaptic release sites by identifying regions of interest (ROI) (Figure 1E) using a series of image pre-processing steps (Figure S1B-E). We found that isoflurane exposure impaired the density and intensity of synaptopHluorin release at these cholinergic synapses (Figure 1F). However, this was not due to changes in the average imaged area (Figure 1G). Rather, the total relative activity (amount of synaptopHluorin release per total release area) and ROI density (the number of ROIs per total neuron size) were significantly decreased under isoflurane anesthesia (Figure 1H-J). In summary, isoflurane impairs evoked cholinergic neurotransmission and reduces the density of release sites in the *Drosophila* brain.

**Figure 1 –.**
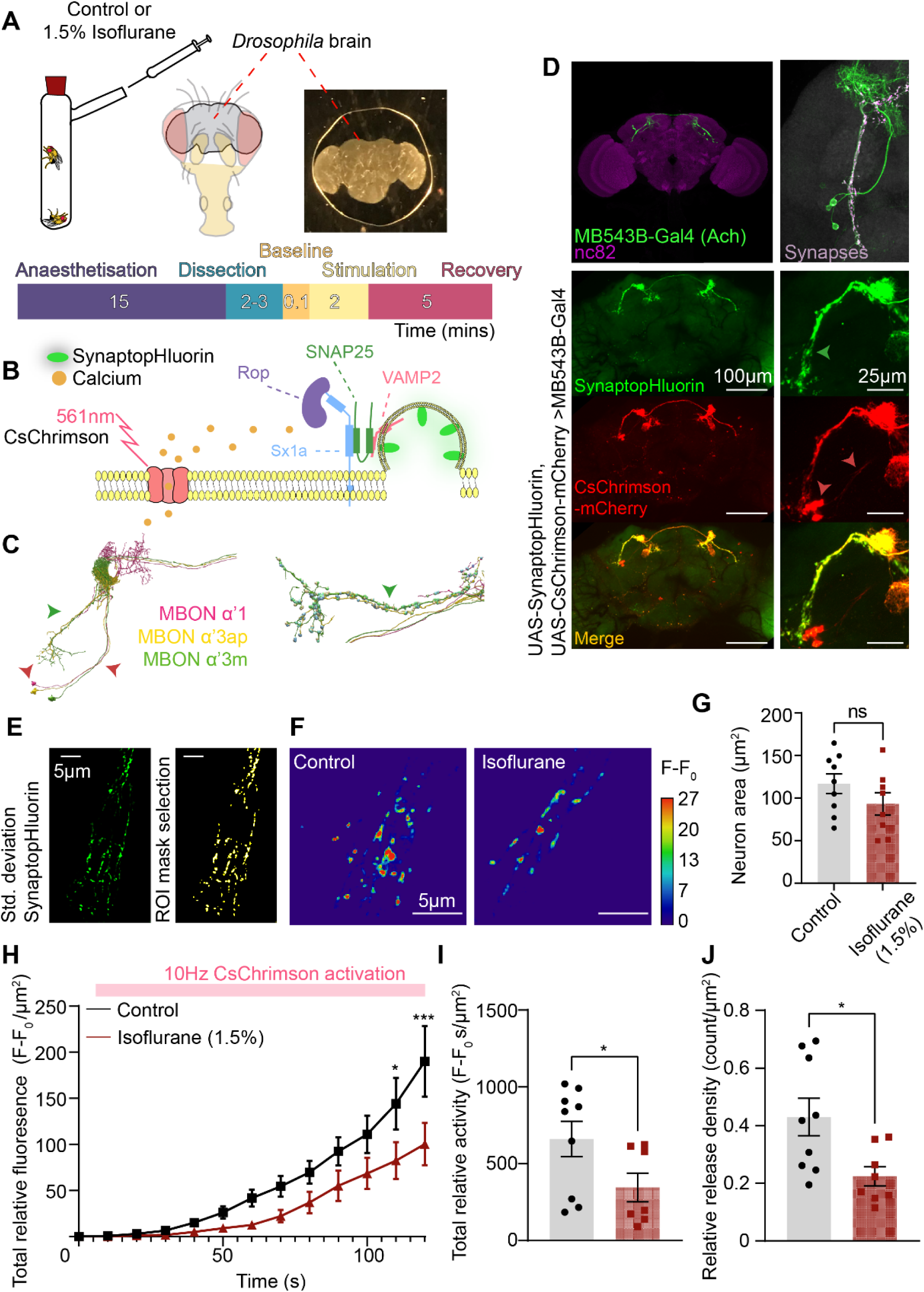
Isoflurane impairs neuroexocytosis at cholinergic synapses in the Drosophila brain. **(A)** Isoflurane anesthesia, equivalent to a 1.5% dose, was induced in female *Drosophila* flies in an air-tight chamber for 15 minutes. The control condition consisted of only air. Brains were then dissected and mounted, and imaged using a standard protocol of 10s baseline, 2min stimulation, and 5min recovery. **(B)** Schema of SynaptopHluorin activity in response to calcium triggered SNARE mediated release via CsChrimson activation. **(C)** Neural skeleton of MB543B-Gal4 derived from FIB-SEM *Drosophila* brain volume (left) with a close up of the synapses (right). Green arrow matches SynaptopHluorin expression in **A**. **(D)** MB543B-Gal4 expression pattern in the *Drosophila* brain immunostained for the pre-synaptic marker bruchpilot (nc82), image taken from Janelia FlyLight split-Gal4 collection. Confocal images of UAS-SynaptopHluorin (top), UAS-CsChrimson-mCherry (middle), and their overlap (bottom) expressed in the cholinergic split-Gal4 (MB543B-Gal4). SynaptopHluorin expression is enriched in synapses (green arrow) but not in cell bodies or extrasynaptic compartments (red arrows). **(E)** Standard deviation Z-projection of the SynaptopHluorin fluorescence change observed during CsChrimson activation (left). A region of interest (ROI) selection is applied to individual release sites for analysis (right). **(F)** Comparison of SynaptopHluorin fluorescence changes under control (left) or isoflurane (right) during CsChrimson activation. Isoflurane decreases the density of release areas and peak fluorescence from baseline. **(G)** The average total neuron area between control and isoflurane conditions was not significantly different (n = 9 control, n = 8 isoflurane, *p* = 0.277, Mann-Whitney test, ± SEM). **(H)** Activity traces of SynaptopHluorin release in control (black) and isoflurane (red) conditions (* *p*<0.05, *** *p*<0.001 Šidák’s multiple comparison test). **(I)** Total relative activity (average change in fluorescence per µm^2^) was significantly decreased under isoflurane anesthesia (*p* = 0.0079, Mann-Whitney test, ± SEM). **(J)** In addition to decreased release, the density of release sites (number of ROIs relative to total neuron area in µm^2^) was significantly decreased under isoflurane anesthesia (*p* = 0.0206, Mann-Whitney test, ± SEM).

Previous research on fly larval motoneurons found that the intravenous general anesthetic propofol differentially impaired transmitter release at active zones of varying size (Karunanithi *et al*., 2020). Propofol halved the total release from all motoneuron synapses, however larger synapses were less impaired than smaller ones due to their greater number of active release sites (Karunanithi *et al*., 2020). To examine whether isoflurane was similarly affecting all release sites in these cholinergic brain synapses, we classified regions of interest (ROI) based on size to observe if there were any differences (Figure 2A). ROIs were defined as either small (<0.08 µm^2^), medium (0.09 – 0.42µm^2^), or large (>0.43µm^2^) based on the k-means clustering of 470 release areas from the control condition (Figure 2A, see Methods). When looking at just the average fluorescence from each release size group for both control and isoflurane conditions, all ROIs show a continuous increase in synaptopHluorin release over time (Figure 2B). However, when looking at the absolute number and relative frequency of each release group, we find that most of the ROIs fall within the small and medium category (Figure 2C). The bulk of the total activity from these cholinergic synapses then are contributed to predominately by release sites <0.42µm^2^ (Figure 2C). Isoflurane does not alter the distribution of different release site sizes, with no significant difference in either the absolute count or relative frequency of small, medium, and large ROIs (Figure 2C). Interestingly, however, the number of small ROIs in the isoflurane condition nearly halved when compared to the control, suggesting many small ROIs were silenced (Figure 2C). Although the average fluorescence did not decrease significantly under isoflurane (Figure S2), knowing that smaller ROIs contributed relatively more to cholinergic release prompted us to investigate if the total relative activity (see Methods) within each synapse size group was diminished. When comparing the total relative activity of each group, we observed a significant reduction in release only from the small and medium groups and no change in the large release sites (Figure 2D). This indicates that disruptions to neurotransmission depend on the size or architecture of a synapse, with the smaller synapses being more vulnerable to isoflurane due fewer active zones (Karunanithi *et al*., 2020) or lower amounts of presynaptic proteins to support them (Ullrich et al., 2015). We next turned our attention to the components of the active zone architecture to investigate presynaptic proteins likely to be key targets of isoflurane.

**Figure 2 –.**
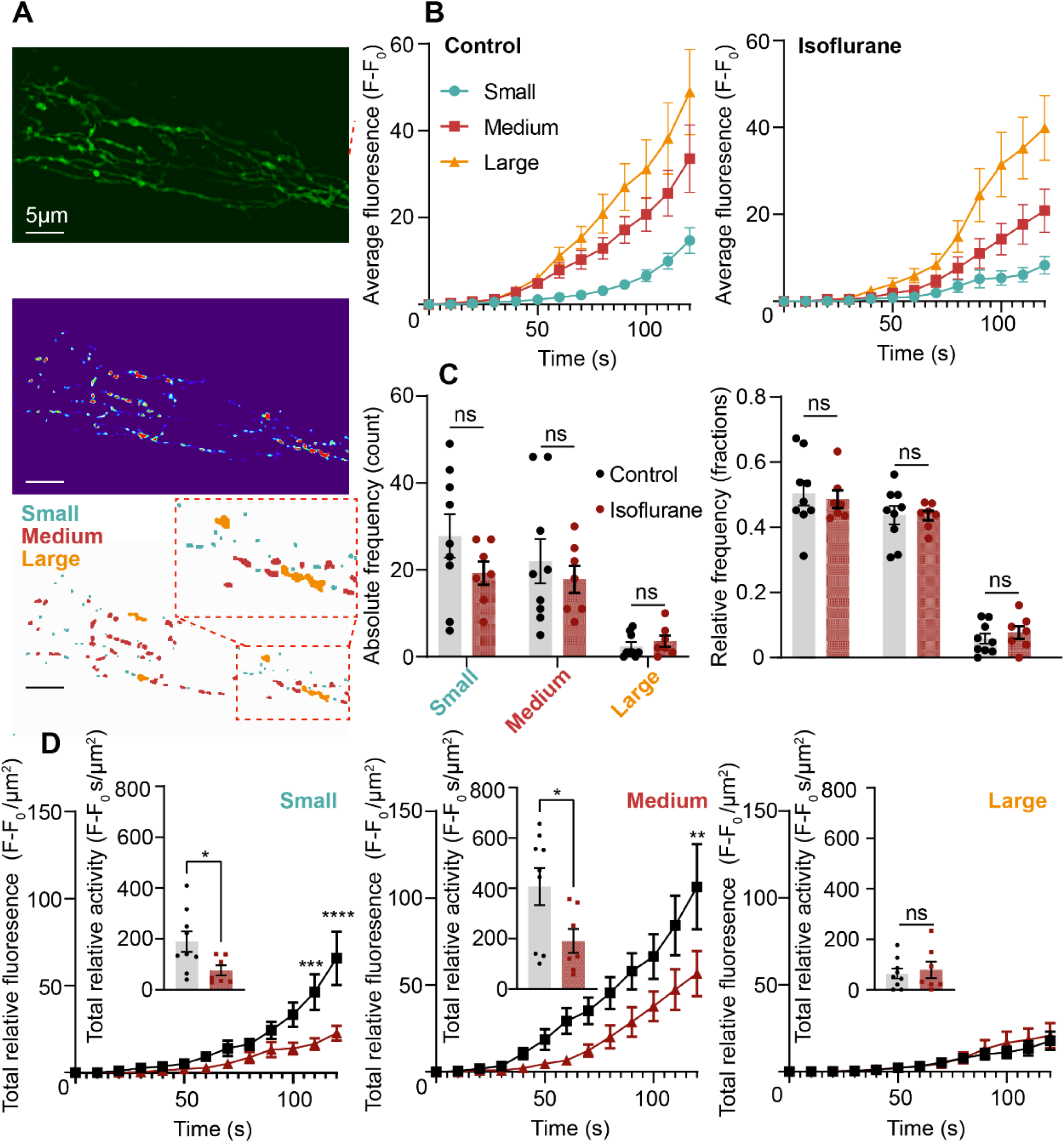
Isoflurane affects smaller, but not larger, active zones by only impairing release. **(A)** (Top) Baseline fluorescence of synaptopHluorin in cholinergic synapses with (middle) the standard deviation of release during CsChrimson activation. (Bottom) Classification of release sites into small (<0.08µm^2^), medium (0.09 – 0.42µm^2^), and large (>0.43µm^2^). Classifications based on *k*-means clustering of individual release areas. (Scale bar 5µm). **(B)** Average fluorescence traces normalised to baseline for control (left) and isoflurane (right) for each of the different release area groups. No significant difference in the activity between control and isoflurane for each group (*p*>0.05 Kruskal-Wallis test, Supp. Figure x). **(C)** Frequency of each release group between control and isoflurane. No difference in the absolute number (left) or the relative frequency (right) of groups within each condition (*p*>0.05 Kruskal-Wallis test, n=9 control, n=7 isoflurane). **(D)** Relative activity for small (left), medium (middle), and large (right) release groups. Only the small and medium release groups showed a significant decrease in total relative fluorescence (** *p*<0.01, *** *p*<0.001, **** *p* < 0.0001 Šidák’s multiple comparison test) and activity (*p*<0.05 Mann-Whitney test).

### Synaptic release machinery proteins are disrupted by isoflurane

General anesthetics disrupt synaptic transmission (Franks, 2008; Franks and Lieb, 1994; Hemmings et al., 2019) by in part by targeting the protein machinery that controls neurotransmitter release (Bademosi *et al*., 2018b; Herring *et al*., 2011; Herring *et al*., 2009; Hines and van Swinderen, 2021; Troup *et al*., 2019; Xie *et al*., 2013; Zalucki *et al*., 2015), including a member of the SNARE complex Sx1a (Bademosi *et al*., 2018b; Hines and van Swinderen, 2021; Nagele et al., 2005). To understand how isoflurane disrupts Sx1a activity in cholinergic synapses that showed impaired neurotransmission, we tagged Sx1a with the photoconvertible fluorophore mEos3.2 and generated transgenic a UAS-Sx1a-mEos3.2 fly strain to perform single-particle tracking and photoactivatable localization microscopy (sptPALM) in our *ex vivo* fly brain preparation (Figure 3A; Supplemental Movie 2). Our Sx1a-mEos3.2 transgene was reliably expressed within the MB543B-Gal4 cholinergic neurons, allowing us to perform sptPALM of photoconverted molecules within the confines of these cholinergic synapses (Figure 3B). To stimulate synaptic release, we opted to use a mVenus-, instead of the mCherry-tagged CsChrimson channelrhodopsin to avoid spectral overlap with photoconverted mEos3.2 signals. To validate our approach and to understand the effects of stimulation on Sx1a-mEos3.2 dynamics, we first compared the mobility and clustering of Sx1a molecules under stimulated or unstimulated conditions (+/- all-trans retinal, see Methods). Upon red light (561 nm) stimulation, Sx1a-mEos3.2 mobility at cholinergic synapses was significantly increased, as quantified by the mean squared displacement (MSD) and a correspondingly increased area under the curve (AUC) (Figure 3C). This indicates that Ca^2+^ influx triggered release increases the mobility and diffusion of these molecules (Padmanabhan et al., 2020). To see if the spatiotemporal clustering of Sx1a-mEos3.2 was altered through CsChrimson activation, trajectories were analysed using segNASTIC software to see if this correlated with changes in molecule diffusion (Figure S3) (Wallis et al., 2021). We revealed that there was a significant decrease in the cluster area of these molecules, but no significant changes to the average lifetime of a cluster or the number of trajectories per cluster (Figure 3D, control black, isoflurane green). A similar increase in the mobility of Sx1a has been previously reported in glutamatergic motor neuron synapses, which could be indicative of the release of Sx1a from presynaptic nanoclusters, binding to other SNARE proteins and the subsequent recruitment to active zones (Bademosi et al., 2016; Padmanabhan *et al*., 2020; Ullrich *et al*., 2015). To confirm that the diffusion and clustering dynamics of Sx1a-mEos3.2 in these cholinergic synapses were physiologically relevant, we impaired synaptic activity during neuronal development with a tetanus toxin light-chain (TeT-LC) (Figure S4). We found that disrupting cholinergic release this way significantly impaired the diffusion of Sx1a molecules, confining Sx1a predominately into clusters (Figure S4). This suggests that Sx1a clustering might be indicative of decreased or impaired neurotransmission.

**Figure 3 –.**
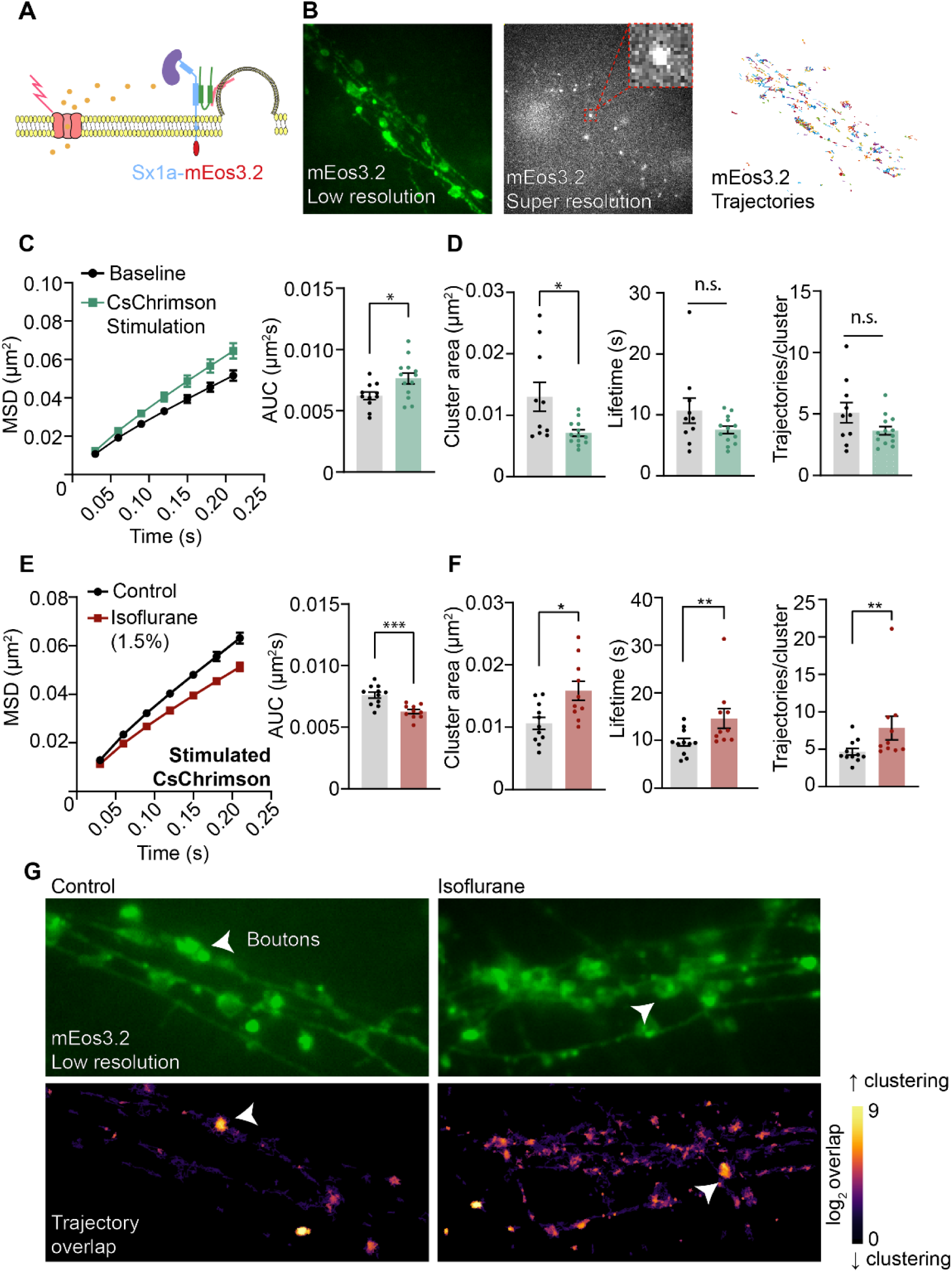
Syntaxin1a mobility dynamics are altered by neuronal Ca^2+^ stimulation and isoflurane anesthesia in cholinergic synapses. **(A)** Schema of the SNARE complex proteins with an mEos3.2 photoconvertible tag attached to the extracellular C-terminus of Syntaxin1a (Sx1a-mEos3.2). **(B)** Low resolution un-photoconverted Sx1a-mEos3.2 in cholinergic synapses (left), super resolution photoconverted Sx1a-mEos3.2 (middle, highlighted inset of a single molecule), and trajectories from analysed Sx1a-mEos3.2 molecules. **(C)** CsChrimson activation of cholinergic synapses increases the mean squared displacement (MSD) and the area under the curve (AUC) of Sx1a-mEos3.2 molecules (n = 11 control, n = 13 isoflurane, *p* = 0.026, Mann-Whitney test, ± SEM). **(D)** Sx1a-mEos3.2 were relieved from clusters under CsChrimson activation (left, *p* = 0.018, Mann-Whitney test) in line with an increase of mobility but did not significantly alter the time spent in clusters (middle, *p* = 0.208, Mann-Whitney test) or the number of trajectories per cluster (right, *p* = 0.208, Mann-Whitney test. All data ± SEM. **(E)** Isoflurane anesthesia significantly decreased the mobility of Sx1a-mEos3.2 in cholinergic synapses in response to CsChrimson activation (n = 11 control, n = 10 isoflurane, *p* = 0.0008, Mann-Whitney test, ± SEM). **(F)** Clustering phenotype of Sx1a-mEos3.2 under isoflurane anesthesia was significantly altered. The cluster area (left, *p* = 0.0197, Mann-Whitney test), lifetime (middle, *p* = 0.0079, Mann-Whitney test), and number of trajectories per cluster (right, *p* = 0.0097, Mann-Whitney test) all significantly increased during CsChrimson activation and isoflurane anesthesia. All data ± SEM. **(G)** Segment overlap plots generated by segNASTIC highlighting increased clustering in the isoflurane condition around boutons (indicated by white arrows).

It has been shown that general anesthetics, both intravenous and volatile, decrease the lateral diffusion of Sx1a on the plasma membrane by trapping them into potentially non-functional nanoclusters (Bademosi *et al*., 2018b; Hines and van Swinderen, 2021). Consistent with these previous findings, we observed a significant decrease in Sx1a-mEos3.2 mobility in activated cholinergic neurons in the presence of isoflurane (Figure 3E). We also found a significant increase in the cluster area, cluster lifetime, and the number of Sx1a trajectories per cluster (Figure 3F). This decrease in mobility and increase in clustering was also found to be activity dependent as we did not observe any effect of isoflurane in unstimulated cholinergic synapses (Figure S5). These data indicate that the trapping of Sx1a on the plasma membrane under isoflurane is not a failure of the stimulation, but rather Sx1a becomes compromised in a way that renders the protein trapped and non-functional (Figure 3G).

To determine if isoflurane acts exclusively on Sx1a, we next measured the effects of isoflurane on the fly homolog of Munc-18a, Rop (Ras opposite), which is responsible for chaperoning Sx1a to the plasma membrane and keeping it in a closed conformation until both interact with other SNARE-associated proteins at active release sites (Dulubova, 1999; Harrison et al., 1994; Rowe et al., 2001). To do this, we generated a UAS-Rop-mEos3.2 transgenic fly line and measured the dynamics of Rob-mEos3.2 mobility at the same cholinergic synapses (Figure 4A-B). Like Sx1a, isoflurane decreased Rop-mEos3.2 lateral diffusion during CsChrimson activation, as evidenced by reductions in both the MSD and AUC (Figure 4C). This was accompanied by a significant increase in the cluster area of Rop molecules but no change in lifetime or trajectories per cluster (Figure 4D). Unlike Sx1a, isoflurane does not cause the trapping of Rop nanoclusters, presumably because it is still able to disassociate from Sx1a upon reaching an active release site. This suggests that a Sx1a/Rop (Sx1a/Munc18) together present a target for isoflurane, also potentially revealing that the diffusion and clustering phenotype observed may be two separate phenomena. Decreases in diffusion could be due some interaction with the Sx1a/Rop complex (Han et al., 2010) whereas clustering of Sx1a could occur post-recruitment to an active zone, without Rop involvement. It is for example understood that Munc18a chaperones Sx1a to the plasma membrane, before both proteins terminate their journey to active zones (Arnold et al., 2017; Kasula et al., 2016; Lai et al., 2017; Shu et al., 2019; Yang et al., 2022).

**Figure 4 –.**
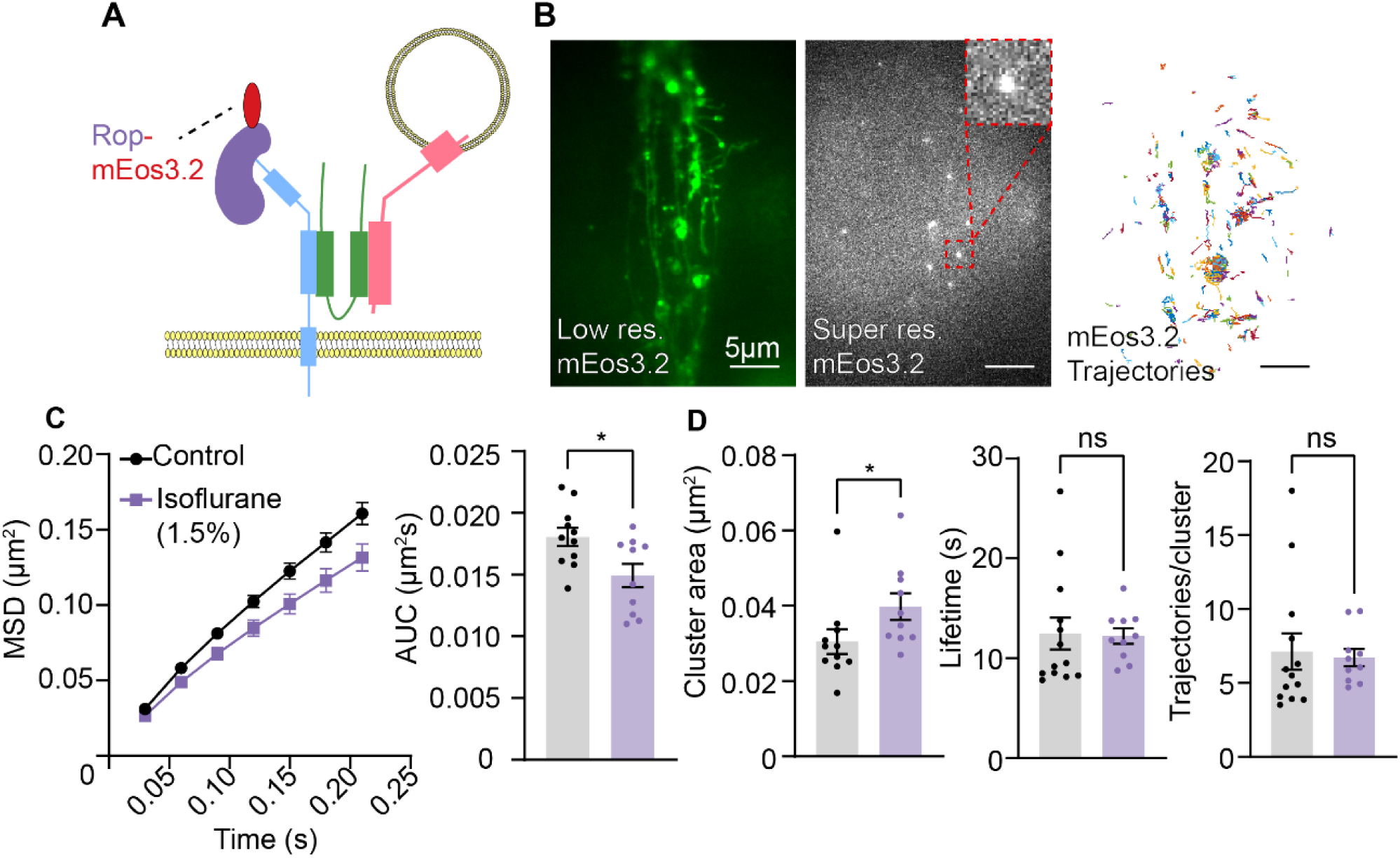
Rop experiences lateral trapping and clustering under isoflurane anesthesia in cholinergic synapses. **(A)** Schema showing the SNARE complex with Rop tagged with an mEos3.2 photoconvertible tag on its C-terminus. **(B)** Low resolution (left), super resolution photoconverted (middle), and trajectories (right) of Munc18-mEos3.2 molecules in cholinergic synapses using the split MB543B-Gal4. **(C)** The MSD and AUC for Munc18-mEos3.2 mobility during CsChrimson activation significantly decreased under isoflurane anesthesia (n = 11 control, n = 10 isoflurane, *p* = 0.0295, Mann-Whitney test, ± SEM). **(D)** In conjunction with a decrease in mobility, Munc18-mEos3.2 molecules experienced a significant increase in the cluster area consistent with Sx1a-mEos3.2 (*p* = 0.0159, Mann-Whitney test, ± SEM).

We have so far found that isoflurane affects the mobility of Sx1a and Rop at pre-synapses, however we still did not know whether the same effects can be observed for Sx1a (or Rop) that is localized in extrasynaptic compartments (Figure 5A) (Maidorn et al., 2019; Ribrault et al., 2011). Interestingly, during CsChrimson activation extrasynaptic Sx1a also displayed a significant decrease in lateral diffusion under isoflurane anesthesia, as measured by the MSD and AUC (Figure 5B). However, in contrast to Sx1a at the synapse, there was no clustering phenotype under isoflurane observed in extrasynaptic regions (Figure 5C). This suggests that while lateral trapping of Sx1a (or Rop) seems to be a common feature of general anesthesia, the nanoclustering phenotype may be dependent on other Sx1a-interacting proteins at the synapse (Figure 5D).

**Figure 5 –.**
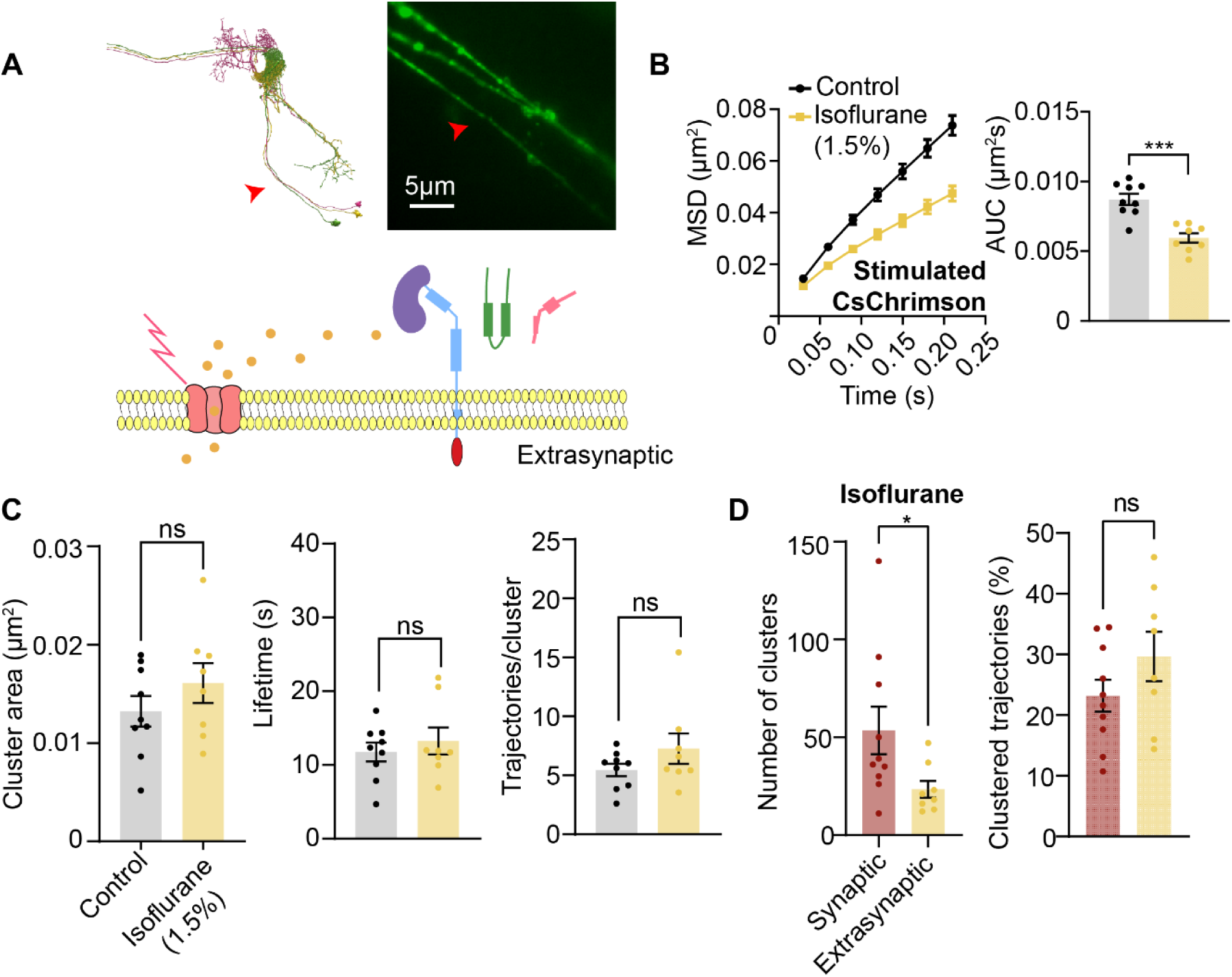
Isoflurane fails to cluster syntaxin1A in extrasynaptic compartments of cholinergic neurons. **(A)** Neural skeleton of MB543B-Gal4 derived from FIB-SEM *Drosophila* brain volume (left) with a close up of the extrasynapses expressing Sx1a-mEos3.2 (right). Red arrow indicates area of close up. **(B)** Isoflurane anesthesia is able to significantly restrict CsChrimson activated Sx1a-mEos3.2 mobility in the absence of synaptic architecture by decreasing the MSD and AUC (n = 9 control, n = 8 isoflurane, *p* = 0.0006, Mann-Whitney test, ± SEM). **(C)** Compared to the synaptic compartment, isoflurane was unable to alter the clustering dynamics of Sx1a-mEos3.2 in the extrasynapse. No significant change in cluster area (left, *p* = 0.37, Mann-Whitney test), lifetime (middle, *p* > 0.999, Mann-Whitney test), or number of trajectories per cluster (right, *p* = 0.541, Mann-Whitney test) was observed. All data ± SEM. **(D)** Comparison between the number of clusters in the presence of isoflurane between synaptic and extrasynaptic compartments was significantly lower for extrasynapses (*p* = 0.043, Mann-Whitney test), however the number of trajectories that were clustered between the two were not significantly different (*p* = 0.203, Mann-Whitney test). All data ± SEM.

### Neurotransmission and Sx1a dynamics in GABAergic and glutamatergic neurons are unaffected by isoflurane

Recent *in vitro* studies suggest that general anesthetics specifically target excitatory but not inhibitory neurotransmitter systems (Koyanagi *et al*., 2019; Speigel et al., 2022; Zhou *et al*., 2019). A possible explanation for this is that the distribution of SNARE protein isoforms is not unilateral across neurotransmitter subtypes (Benagiano et al., 2011). In addition to VAMP2, the third partner involved in the formation of SNARE complexes is SNAP25 (Südhof, 2012). Previous work has shown that botulinum neurotoxin E, which cleaves SNAP25 irreversibly, alleviates the effects of propofol on Sx1a dynamics in cultured PC12 cells (Bademosi *et al*., 2018b). It has been suggested that subpopulations of Gad1-positive neurons in rats and humans lack SNAP25 immunoreactivity, despite the presence of *SNAP25* mRNA (Benagiano *et al*., 2011; Davie et al., 2018; Frassoni et al., 2005; Garbelli et al., 2008; Mandolesi et al., 2009; Verderio et al., 2004). SNAP25 immunoreactivity has also been shown to be absent from some mushroom body neuropils in the fly brain, with the suggestion that the SNAP24 isoform replaces SNAP25 expression in these neurons to facilitate vesicle fusion (Niemeyer and Schwarz, 2000; Vilinsky et al., 2002). To see if inhibitory MBONs in the fly brain lack SNAP25 expression, we selected a GABAergic and glutamatergic split-Gal4 drivers (Figure 6A) and performed immunohistochemistry for SNAP25 on the brains of both inhibitory neurotransmitter subtypes as well as our excitatory cholinergic circuit (Figure 6B). The role of glutamate in the fly CNS remains ambiguous (Chvilicek et al., 2020) and has been shown to exert inhibitory functions, whereas it is known to be excitatory at the neuromuscular junction (Jan and Jan, 1976; Liu and Wilson, 2013; Molina-Obando et al., 2019). The SNAP25 antibody stained the entire brain and was detected with an Alexa-647 fluorescent probe. We found that for the cholinergic (MB543B-Gal4) neurons there was a large amount of overlap between the mVenus expression in the neurons and SNAP25 antibody staining, whereas the GABAergic (MB112C-Gal4) and glutamatergic (MB433B-Gal4) neurons displayed no significant overlap (Figure 6B; Figure S6A-C). The SNAP25 immunoreactivity in GABAergic and glutamatergic neurons was predominately found around the periphery of their dendritic arbours (Figure 6B, right).

**Figure 6 –.**
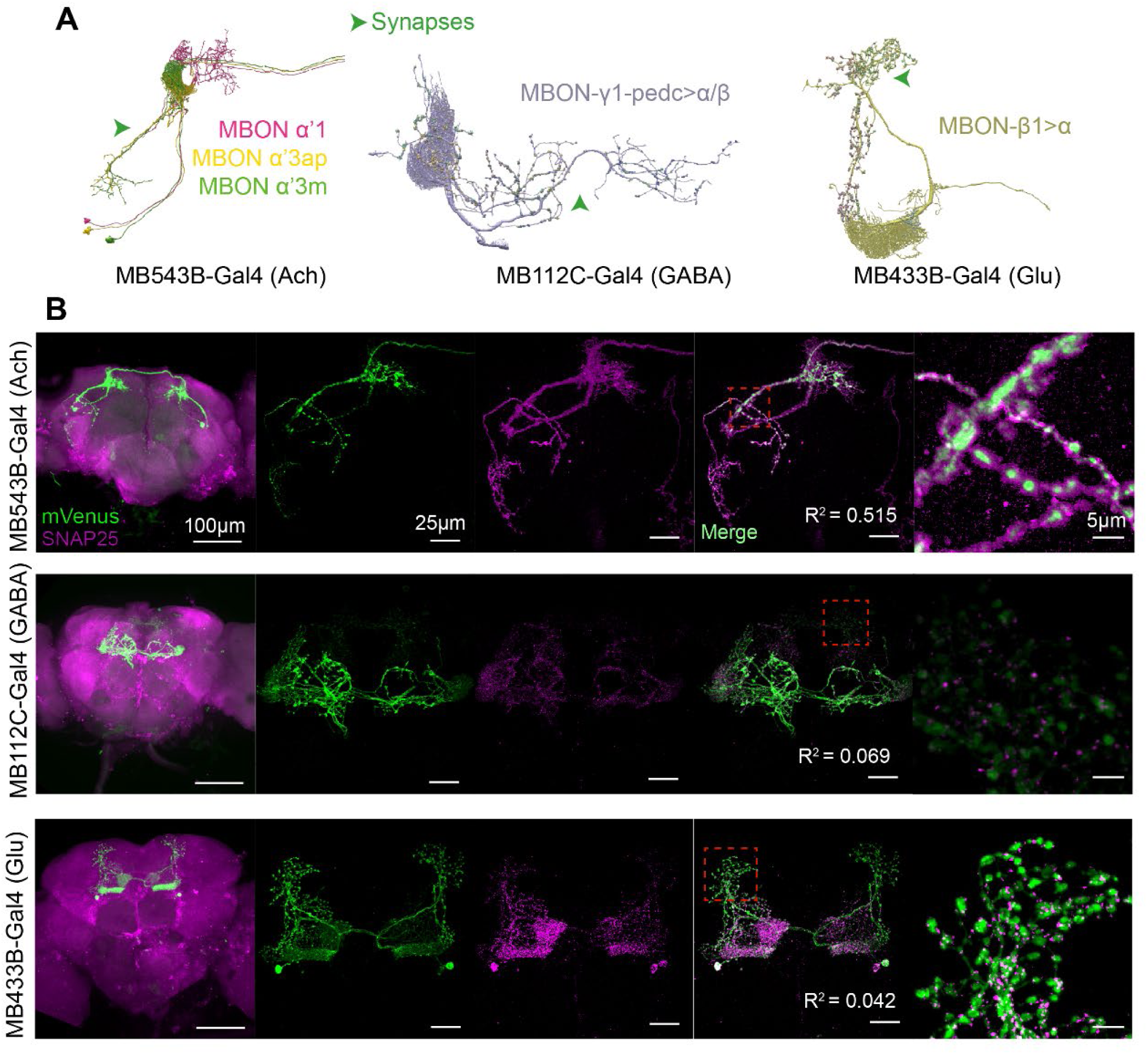
Inhibitory MBON neurons lack SNAP25 expression. **(A)** Neural skeletons of MB543B-Gal4 (Ach., left), MB112C-Gal4 (GABA, middle), and MB433B-Gal4 (Glu., right) from the *Drosophila* FIB-SEM connectome. Green arrows indicate location of synapses. (B) Co-localisation of SNAP25 immunostaining to each neurotransmitter system for MB543B-Gal4 (top), MB112C-Gal4 (middle), and MB433B-Gal4 (bottom). SNAP25 expression is enriched in the excitatory cholinergic synapses (R^2^ = 0.515) whereas in the inhibitory GABAergic and glutamatergic neurons there is no detectable puncta within synapses (R^2^ = 0.069 for GABA, R^2^ = 0.042 for glutamate).

We next performed synaptopHluorin imaging in synapses of the GABAergic (MB112C-Gal4) and glutamatergic (MB433B-Gal4) neurons, to determine if isoflurane also impaired release from these inhibitory neurons (Figure 7A,D) (Aso *et al*., 2014). For both GABA and glutamate, no significant difference was observed for the total relative fluorescence, activity, or ROI density of synaptopHluorin release in the presence of isoflurane when compared to control (Figure 7B,E). Between release size groups, isoflurane did not change the release characteristics of the different ROIs, however there was a significant decrease in the relative frequency of the small release group for glutamate (Figure 7C,F). Compared with the cholinergic neurons, the larger ROIs contributed more of the total relative activity in GABAergic and glutamatergic neurons (Figure 7G,H). Since a larger proportion of release from cholinergic synapses was from small and medium sized ROIs, resistance to the overall decreased transmitter release in inhibitory synapses could be due to the comparatively lower activity in smaller ROIs (Figure 7G,H).

**Figure 7 –.**
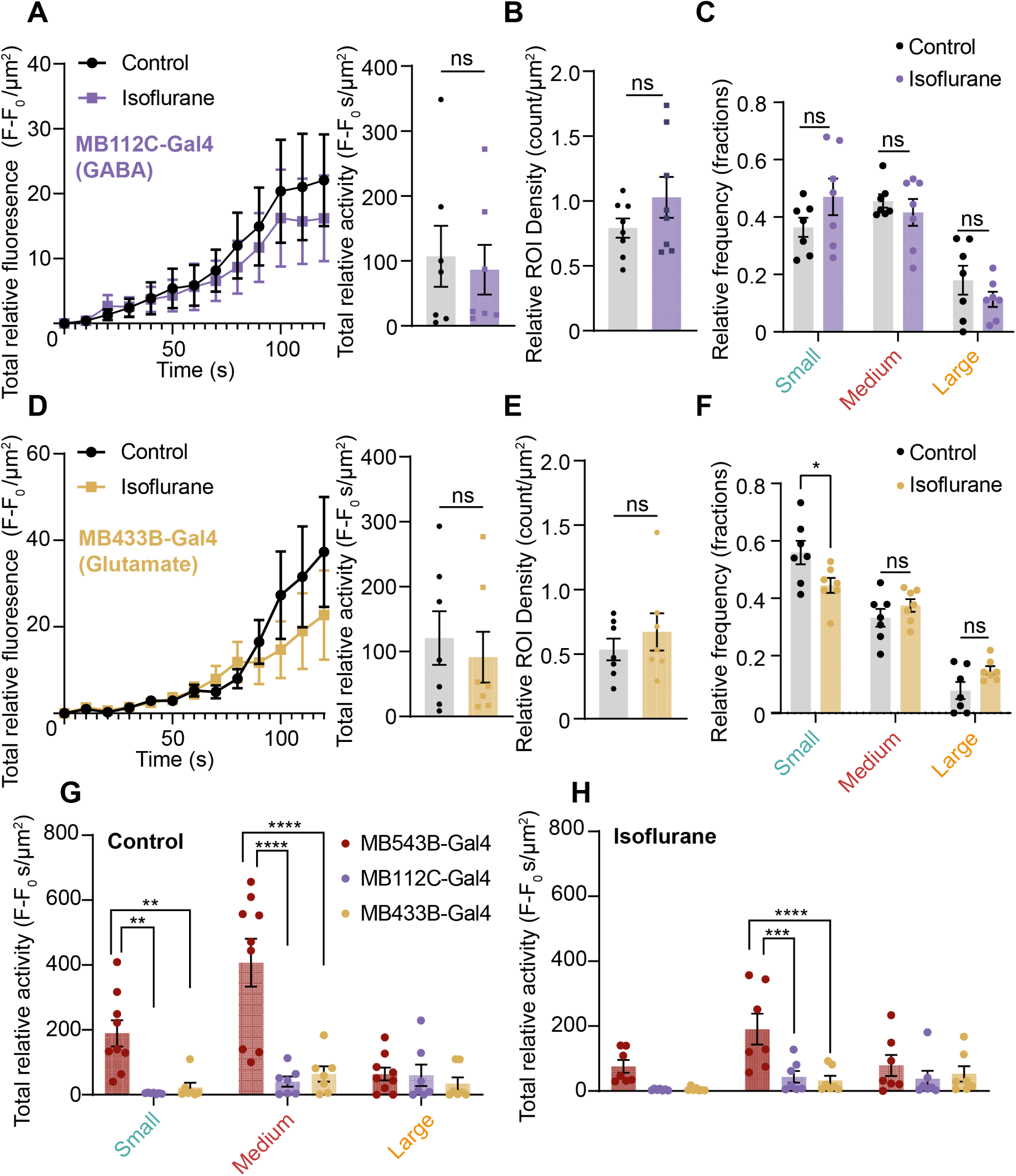
Synaptic release in inhibitory MBONs is intact with isoflurane. **(A)** (Top) UAS-SynaptopHluorin expressed into GABAergic synapses with the (bottom) F-F_0_ of release during a 2-minute CsChrimson stimulation. **(B)** Total relative fluorescence and activity from GABAergic synapses is unaffected by isoflurane, with no change in the relative ROI density (*p*>0.05 Mann-Whitney test, n = 7 control & isoflurane). **(C)** No change to the relative distribution of different release site sizes, however compared to cholinergic synapses the bulk of the total relative activity is contributed by large release sites (*p*>0.05, Šidák’s multiple comparison test). **(D)** (Top) UAS-SynaptopHluorin expressed into glutamatergic synapses with the (bottom) F-F_0_ of release during a 2-minute CsChrimson stimulation. **(E)** Like GABA synapses, release from glutamatergic synapses is also unaffected by isoflurane anesthesia (*p*>0.05 Mann-Whitney test, n = 7 control & isoflurane). **(F)** Small release sites significantly decreased in the isoflurane condition (* *p*<0.05, Šidák’s multiple comparison test), however there was no difference in the amount of release per region size (*p*>0.05, Šidák’s multiple comparison test). **(G)** Comparison of total relative activities for the control condition between small, medium, and large for the 3 neurotransmitter circuits. MB543B-Gal4 had significantly higher activity from small and medium release sites than MB112C-Gal4 and MB433B-Gal4 (** *p*<0.01, **** *p*<0.0001, Šidák’s multiple comparison test) whereas the large release sites were approximately the same. **(H)** Same comparison as (G) but for the isoflurane condition. Only the medium release group for MB543B-Gal4 was significantly higher than MB112C-Gal4 or MB433B-Gal4 (*8* *p*<0.001, **** *p*<0.0001, Šidák’s multiple comparison test).

We questioned whether resistance to isoflurane in inhibitory neurons was solely evident in our synaptopHluorin readouts, or whether sx1a dynamics were also resistant to the drug in these neurons, To address this, we expressed Sx1a-mEos3.2 in either set of neurons (Figure 8A,C). For both GABAergic and glutamatergic synapses exposed to isoflurane, we observed no significant decrease in mobility and no increase in clustering for Sx1a molecules (Figure 8B,D). This suggests that the isoflurane resistance we observed at the level of exocytosis may result in part from resistance effects already present in the presynaptic release machinery of these inhibitory neurons. One obvious difference with cholinergic neurons, as discussed above, is the absence of SNAP25.

**Figure 8 –.**
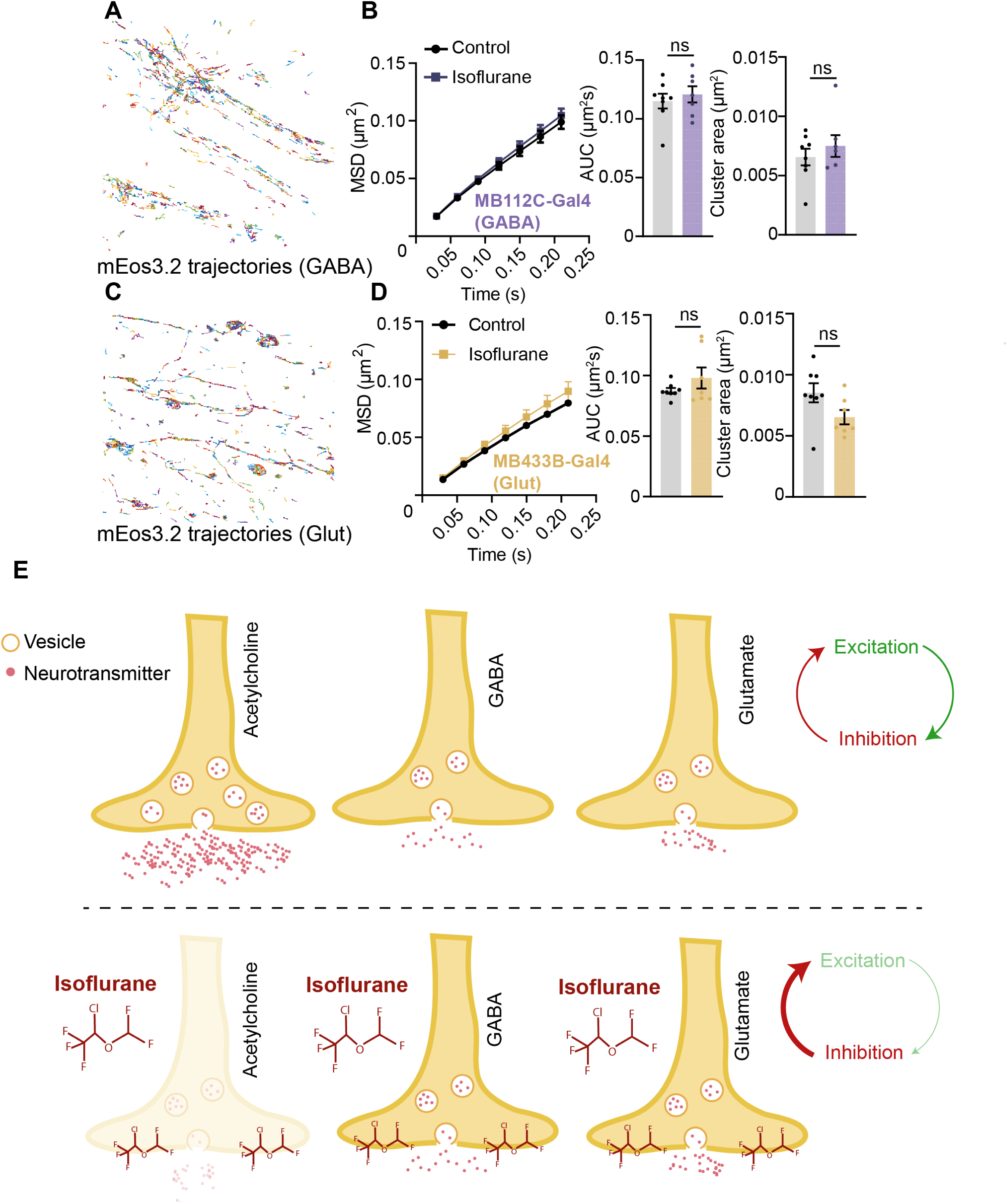
Syntaxin1a dynamics are unaffected by isoflurane in SNAP25 deficient synapses. **(A)** Sx1a-mEos3.2 trajectories for MB112C-Gal4 (GABA) inhibitory synapses. **(B)** Sx1a-mEos3.2 MSD, AUC, and cluster area is unaffected by isoflurane anesthesia in GABA synapses (*p*>0.05 Mann-Whitney test, n = 7 control & isoflurane). **(C)** Sx1a-mEos3.2 trajectories for MB433B-Gal4 (Glut) inhibitory synapses. **(D)** Sx1a-mEos3.2 MSD, AUC, and cluster area is unaffected by isoflurane anesthesia in glutamatergic inhibitory synapses (*p*>0.05 Mann-Whitney test, n = 7 control & isoflurane). **(E)** Schema of the effects of isoflurane on excitatory vs inhibitory synapses. In the presence of isoflurane, only excitatory neurotransmitter release is impaired. The brain under general anesthesia in concert with post-synaptic activation of GABA receptors experiences successive bouts of inhibition by reducing the overall excitatory potential.

## Discussion

Over the last few decades, several potential targets of general anesthetics in the CNS have emerged to help explain these drugs’ unique ability to produce a rapid loss of consciousness and unresponsiveness (Franks, 2008; Hemmings *et al*., 2019). Explanations for general anesthesia have largely focussed on mechanisms that reduce neuronal excitability through post-synaptic potentiation of inhibitory receptors such as GABA_A_ (Franks and Lieb, 1984; 1994; Tomlin et al., 1998). Recently, a complementary set of potential mechanisms have begun to emerge by studies examining presynaptic disruptions in neurotransmission (Baumgart *et al*., 2015; Franks, 2008; Hemmings *et al*., 2019; Hemmings et al., 2005; Herring *et al*., 2011; Herring *et al*., 2009; Koyanagi *et al*., 2019; Ouyang et al., 2003; Speigel and Hemmings, 2021; Torturo *et al*., 2019; van Swinderen and Kottler, 2014; Xie *et al*., 2013; Zhou *et al*., 2019). However, it remains unclear which of the presynaptic release processes are relevant to anesthesia endpoints, and in which neurons specifically. Our study uncovers unequal effects of isoflurane on different neurotransmitter systems, defined by whether they are excitatory or inhibitory, and by whether they express the SNARE protein SNAP25. This presents a presynaptic mechanism for general anesthesia centred on impaired excitatory neurotransmission, that complements postsynaptic inhibitory processes.

It is now well established that there are presynaptic effects of general anesthetics that reduce the reliability of chemical neurotransmission (Bademosi *et al*., 2018b; Baumgart *et al*., 2015; Hemmings *et al*., 2005; Herring *et al*., 2011; Herring *et al*., 2009; Karunanithi *et al*., 2020; Koyanagi *et al*., 2019; Speigel and Hemmings, 2021; Torturo *et al*., 2019; Zalucki *et al*., 2015), thereby impairing coordinated neural processes across the brain (Liu et al., 2013; Ranft et al., 2016). Our growing understanding of the control of neurotransmitter release highlights its multifaceted and complicated nature and is helping to reveal general anesthetic effects on synaptic transmission (Südhof, 2012; Sudhof and Rizo, 2011; Sudhof and Rothman, 2009). SNARE-associated molecules (Sx1a, SNAP25, VAMP2, Munc18) drive vesicular fusion with the plasma membrane via a Ca^2+^ triggered mechanism that requires the propagation of action potentials in most animals (Südhof, 2012). We found that CsChrimson stimulated synaptic release is impaired using our synaptopHluorin readout in excitatory cholinergic synapses (Figure 1H-J), but not in the inhibitory GABAergic or glutamatergic neurons (Figure 7B & E). This effect correlates with Sx1a lateral diffusion and clustering readouts at the synapses of these distinct neurons (Figure 3E-F, 5B-C, 8B,D). It has been suggested that differences in expression of voltage gated Ca^2+^ channels are responsible for the differential sensitivity to isoflurane in different neuronal subtypes (Baumgart *et al*., 2015; Torturo *et al*., 2019). Our results suggest that the role of Ca^2+^ channels might not be directly relevant, at least in the *Drosophila* brain, because we expect that the amounts of Ca^2+^ entry induced by the activation of CsChrimson are comparable in all three circuits tested. It is therefore reasonable to conclude that the effects we observed on the diffusion and clustering of Sx1a must be independent of Ca^2+^ channel activity. This provides further evidence that general anesthetics such as isoflurane and propofol act downstream from presynaptic ion channels (Wang *et al*., 2020) by interacting directly with SNARE molecules (Bademosi *et al*., 2018b; Hines and van Swinderen, 2021; Nagele *et al*., 2005). However, it remains likely that diverse presynaptic targets work synergistically to impair neurotransmission (Hemmings *et al*., 2019).

How isoflurane mediated Sx1a lateral trapping and clustering might cause decreased neurotransmission remains unclear, particularly since the precise location of clustering relative to active release sites is unknown. A previous study focussing on the intravenous anesthetic propofol proposed that Sx1a molecules are trapped by the drug into non-functional nanoclusters on the plasma membrane, prior to SNARE formation at the active zone (Bademosi *et al*., 2018b). From our results we can conclude that whilst lateral diffusion for Sx1a decreases in both extrasynaptic and synaptic compartments, clustering only occurs at synapses (compare Figure 3F and Figure 5C). This indicates that synaptic release architecture must likely be present for Sx1a molecules to cluster under isoflurane anesthesia, and that increased clustering potentially occurs after Sx1a recruitment to active zones (Figure 3G). This conclusion is consistent with the apparent decrease in lateral diffusion and increased clustering of Rop (Munc18, Figure 4C-D), likely due to its strong interaction with Sx1a immediately prior to SNARE formation (Burkhardt et al., 2008).

Considered together with previous findings, our comparison of different neurotransmitter systems in the fly brain provides an explanation for why certain synapses were affected by isoflurane but not others. Earlier work using propofol showed that SNAP25 is required for the general anesthetic to be able to impair Sx1a dynamics: cleaving SNAP25 with a botulinum neurotoxin E light chain reversed the lateral trapping effect of propofol on Sx1a and resulted in an increase in Sx1a diffusion, and this effect was replicated using a deletion mutant of Sx1a that abrogates SNAP25 interaction (Bademosi *et al*., 2018b). We observed a significant reduction in Sx1a mobility in both synaptic and extrasynaptic compartments of a cholinergic circuit, likely due to the fact that SNAP25 is present in both areas (Maidorn *et al*., 2019). In contrast, Sx1a lateral diffusion and neuroexocytosis was intact in GABAergic and glutamatergic neurons under isoflurane anesthesia (Figures 7 & 8). This suggests an important difference in release machinery components between these different neuron types. Indeed, neuronal circuits can differentially express particular isoforms of SNAREs (Augustin et al., 1999; Kádková et al., 2019; Ruiz-Montasell et al., 1996), especially SNAP25 where it has been shown to be absent in inhibitory neurons (Frassoni *et al*., 2005; Garbelli *et al*., 2008; Holt et al., 2006; Mandolesi *et al*., 2009; Su et al., 2001; Verderio *et al*., 2004). Given that we saw a clear isoflurane phenotype in the excitatory cholinergic system and no effect at all in the inhibitory GABAergic and glutamatergic circuits, it was reasonable to hypothesise that the latter neurons may lack SNAP25 expression and is exactly what we found (Figure 6B). Together, these results present SNAP25 as a possible target for general anesthetics, probably as an interaction with Sx1a / munc-18. Indeed, previous studies have detected direct interactions of general anesthetics to Sx1a and SNAP25 proteins independently (Nagele *et al*., 2005; Weiser et al., 2013). One possibility is that a different isoform of SNAP25 driving release in inhibitory neurons is less sensitive to general anesthetics. For example, SNAP24, a mammalian SNAP23 or SNAP25b ortholog in *Drosophila*, has been shown to be capable of driving chemical neurotransmission in mushroom body neurons lacking SNAP25 immunoreactivity (Niemeyer and Schwarz, 2000; Vilinsky *et al*., 2002).

In conjunction with a potential SNAP25 protein explanation for insensitivity to general anesthetics, another interesting observation from our study is the resistance that larger release sites experience during isoflurane anesthesia (Figure 2D). Amongst the three neurotransmitter systems we tested, the relative contribution of each size group to the total activity skewed toward larger ROIs in inhibitory MBONs (Figure 7G,H). This is likely related to the way some inhibitory synapses develop, typically less efficient at vesicle recycling and driven by more tonic release mechanisms (Bae et al., 2020; Häusser and Clark, 1997). Studies have shown a correlation between the size and release capacity of an active zone and the number of tethered vesicles by SNARE molecules localised to them, which could provide resistance to the effects of general anesthetics due to the higher amount of release architecture present (Han et al., 2017). However, release site size alone could not fully account for differences between neurotransmitter groups as there was no direct effect on the diffusion dynamics of Sx1a molecules in inhibitory synapses (Figure 8B,D). These findings indicate potentially two different presynaptic mechanisms: 1. Pre-synaptic targeting of excitatory neurons by severely disrupting the smaller and more active release sites, and 2. Specific targeting of Sx1a molecules and cognate SNARE partners (including Munc18, Figure 4), which have previously been reported to be binding targets of several general anesthetics (Nagele *et al*., 2005; Woll et al., 2016; Xie *et al*., 2013). It is also likely that these presynaptic effects are not independent.

From our findings, it is evident that not all neurons in the fly CNS are affected by general anesthetics in the same way and that there are clear differences between neurotransmitter subtypes. Our findings align well with other studies in mammals where excitatory glutamatergic neurons show decreased neurotransmission in response to isoflurane anesthesia, but not GABAergic neurons (Koyanagi *et al*., 2019; Speigel and Hemmings, 2021; Torturo *et al*., 2019). Considering our findings together with postsynaptic theories, we speculate that the brain under general anesthesia is potentially experiencing successive bouts of inhibition to cause a loss of consciousness and responsiveness: first a circuit-specific inhibition of excitatory activity by potentiation of postsynaptic GABA_A_ receptors embedded in arousal systems, followed by a more global downscaling of presynaptic release from excitatory neurons (Figure 8E) (van Swinderen and Kottler, 2014). The relative contribution to different anesthesia endpoints as well as recovery dynamics should be discernible using neuron-specific expression systems in behaving animals.

## Methods

### Transgenic fly strains

UAS-Syntaxin1A-mEos3.2 and UAS-Rop-mEos3.2 constructs for super resolution microscopy were generated as previously described (Bademosi et al., 2018a). In brief, Drosophila Syntaxin1a and Rop had an mEos3.2 fluorophore attached to it by replacing the stop codon for a linker with the sequence gaggtaccgcgggcccgggatccaccg. Synthesis and cloning of this construct were performed by VectorBuilder™ and was cloned into a 5xUAS Hsp70 Drosophila expression vector with a mini-white gene. PhiC31 integration into the Drosophila genome was performed by Genetivision Corporation™ into the attP40 landing site on the second chromosome.

### Fly stocks

All flies used in the study were reared on a standard yeast agar food at 25°C with a 12-hour day/night cycle. Flies used for the CsChrimson stimulation were reared on the same food supplemented with 0.2mM all-trains retinal (ATR) for at least 72 hours prior to imaging. w1118 UAS-Syntaxin1A-mEos3.2/Cyo; Sb/Tm6Tb and w1118 UAS-Rop-mEos3.2/Cyo; Sb/Tm6Tb stocks were combined with w1118 Sco/Cyo; UAS-CsChrimson-mVenus/Tm6Tb to generate the working strain w1118 UAS-Syntaxin1A-mEos3.2/Cyo; UAS-CsChrimson-mVenus/Tm6Tb and w1118 UAS-Rop-mEos3.2/Cyo; UAS-CsChrimson-mVenus/Tm6Tb to cross to split-Gal4 drivers. Mushroom body output neuron (MBON) split-Gal4 lines have been previously described (Aso *et al*., 2014) and were ordered from the Bloomington Drosophila Stock Centre (BDSC). For all sptPALM experiments after expressing the working strain into a split-Gal4, female flies were sorted upon eclosing by brief anesthetization on a CO2 pad and imaged 3-5 days later. Females were chosen to control for sexual dimorphisms in expression patterns.

### Adult Drosophila brain dissection and mounting

3-5 day old adult Drosophila brains were prepared for imaging similarly as previously described (Hines and van Swinderen, 2021). For baseline to CsChrimson activation experiments, flies were cold anesthetized by placing them in a glass, air-tight chamber on ice for 30 seconds, then dissected on ice cold PBS and imaged immediately. For all anesthesia experiments flies were placed in a glass, air-tight chamber and exposed to either 5µL air (control) or 100% isoflurane (equivalent to a 1.5% dose) for 15 minutes prior to cold anesthetization and dissection, as determined by gas chromatography (Zalucki *et al*., 2015). For the isoflurane condition, the 15 minutes started after the flies became fully anesthetized (i.e. were behaviourally unresponsive to mechanical stimulation at the base of the glass chamber), which on average took approximately 10-30 seconds per fly.

The imaging buffer was a modified hemolymph-like 3.1 (HL3.1) solution which consisted of 70mM NaCl, 5mM KCl, 1.5 mM CaCl_2_, 2mM MgCl_2_, 5mM HEPES, 115mM sucrose, 5mM trehalose, and pH adjusted to 7.2 with NaHCO_3_ (Sigma-Aldrich). Imaging buffer was made in batches, aliquoted, and stored at −80°C. A fresh aliquot of HL3.1 was defrosted for each day of experiments. Dissected brains are placed in 2µL of imaging buffer on a glass slide and sealed underneath a no. 1.5 22×22mm coverglass rimmed with vacuum grease and nail polish.

### Live cell SynaptopHluorin imaging and processing

Prior to imaging, we bleached the baseline synaptopHluorin fluorescence using 25% power of an 18.72mW 488nm laser at 30msec exposure for 1 minute (Figure S1A). This is because there is a significantly high background fluorescence of synaptopHluorin in *Drosophila* nervous tissue, potentially due to a higher internal vesicle pH compared to mammalian systems. After the photobleaching, the exposure time was set to 100msec and 488nm laser power reduced to 4% to capture synaptopHluorin release. 10 seconds of baseline was captured prior to 2 minutes of 10Hz red light stimulation using 25% power of a 20.6mW 568nm laser. Recovery was captured for 5 minutes after red light stimulation (Figure S1E).

To analyse the data, several pre-processing steps to the raw imaging data were applied to resolve the true delta fluorescence change during stimulation (Figure S1B). Firstly, recordings were truncated to just include the baseline and stimulation frames to allow for more accurate motion correction. The truncates recordings were motion corrected using ImageJ’s ‘Correct 3D Drift’ plugin (Parslow et al., 2014) with ‘Correcting only x & y’, ‘Multi time scale computation’, and ‘Sub pixel drift correction’ all selected. All other settings were left to default. Motion corrected recordings were background subtracted to a rolling ball radius of 5 pixels (0.5 microns). To eliminate baseline fluorescence, a maximum Z-projection of the first 100 frames (baseline prior to red light activation) was generated and used to subtract all the pixel values for the entire recording. Finally, a despeckle noise remover was applied to eliminate background signal incurred due to bleed through of red light into the camera detector. To select ROIs to analyse, a standard deviation Z-projection of the processed recording was generated, and a ‘Li’ auto-threshold was calculated to create a mask and selection of the regions with high increases of fluorescence (Figure S1C). The selection was then applied to the recording and a multi-measure of the mean pixel grey value and area for each individual ROI was generated and analysed in Python. These pre-processing steps were crucial to see the true change in fluorescence throughout stimulation (Figure S1D).

### Quantification of SynaptopHluorin release

Mean grey values and areas measured in each ROI for SynaptopHluorin release were imported into a custom Python script for analysis (available at github.com/AdamDHines/synaptopHluorin-analysis). The fluorescence (F-F_0_) for each ROI of a recording was summed together and divided by the total area of release for the ‘Total relative fluorescence’. The data was down sampled by averaging the total relative fluorescence into 10 second bins over the course of the 2-minute stimulation period. To calculate a proxy for quantal release, a trapezoidal area under the curve (AUC) was calculated for each of the total relative fluorescence curves for the ‘Total relative activity’. The relative ROI density was calculated by taking the number of detected ROIs and dividing it by the total area of the imaged neuron. This was calculated by taking an image of the neuron using 488nm excitation, background subtracting with a 0.5 micron rolling ball radius, and applying a ‘Li’ auto-threshold to create a mask around the neuron (Li and Tam, 1998). This mask was converted to a selection and the total area was calculated. Release area groups were calculated using the *k-*means clustering of 470 ROIs from the control acetylcholine recordings. A maximum of 10 iterations and 3 groups were defined, which repeatedly output the same cluster centroids. The centroids were then used to discretise the ROIs for all synaptopHluorin recordings into small, medium, or large groups and analysed the same way as the total activity.

### Drosophila brain PALM

The method for performing sptPALM in adult fly brains has been extensively described (Hines and van Swinderen, 2021). All super resolution microscopy was done on a Zeiss ELYRA PS.1 to allow simultaneous 405nm and 561nm wavelength laser operation microscope through an Apochromat 100×1.4 NA oil immersion objective. All recordings were captured on an iXon EMCCD camera. Regions of interest were navigated to with a 10x 0.45 NA Plan-Apochromat objective using a 488nm wavelength laser before switching to 100x oil immersion. Brains were illuminated in a highly inclined and laminated optical (HILO) sheet imaging mode with an angle of 47.3° to improve the signal-to-noise ratio. 0.02 – 0.08% power of a 0.6mW 405nm laser was used to photoconvert mEos3.2 from green to red and 25% power of a 20.6mW 561nm laser was used to excite photoconverted mEos3.2 molecules. A Zeiss incubation chamber with a TempModule S attached to it regulated the temperature at 25°C for all imaging experiments. 32,000 images were captured at a 30msec exposure time (the lowest possible for the camera to record at) per recording, with approximately 1-2msec between each frame.

### Single particle tracking of PALM data

Molecule localisation and tracking were performed in the custom MATLAB GUI *Single Particle Analysis* (SPA) which employs the free ImageJ plugin TrackMate (Hines and van Swinderen, 2021; Tinevez et al., 2017). Single Sx1a-mEos3.2 and Rop-mEos3.2 molecules were detected and localised using a Laplacian of Gaussian detection algorithm (Equation 1) with a spot radius of 0.4µm and a variable threshold depending on the signal-to-noise ratio (manually determined). Detection was improved by using median filtering and subpixel localisation. We achieved a localisation precision of approximately 25nm with a point-spread function half-width of 134nm (Figure S3A).

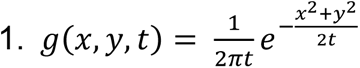

Where x and y are the spatial coordinates and t is the frame time. Molecule tracking between frames was performed by a linear assignment problem (LAP) algorithm (Jaqaman et al., 2008). To be included for analysis, a track had to have a minimum 6 spots and a maximum of 1,000 per trajectory.

Mean squared displacement (MSD) and clustering analysis was all performed using segNASTIC software (Wallis *et al*., 2021). For accurate spatiotemporal clustering analysis, drift correction was applied to the x,y coordinates of trajectories by feeding drift tables calculated in Zeiss ZEN PALM calculated by a model-based drift algorithm (Figure S3B-E). Drift correction did not change the mobility data but did give different results for spatiotemporal clustering indicating the importance of performing drift correction in this analysis (Figure S3F). The area under the curve (AUC) was calculated in Graphpad Prism 9 with a baseline starting at Y = 0, ignoring peaks that are <10% of the distance from minimum to maximum Y, and defining that all peaks must go above baseline. Outliers in MSD data were determined by excluding results that were ± 2 times the standard deviation of the mean area under the curve for each condition. Outliers for clustering data were determined by ROUT method with a maximum false discovery rate of 1% calculated in Graphpad Prism 9.

### Immunohistochemistry of *Drosophila* brains

3–5-day old flies expressing CsChrimson-mVenus in cholinergic (MB543B-Gal4), GABAergic (MB112C-Gal4), or glutamatergic (MB433B-Gal4) neurons were briefly anesthetized on a CO_2_ pad and brains dissected in ice cold PBS. The brains were then fixed in a 4% paraformaldehyde (PFA) solution for 30 minutes at room temperature with rotation to chemically fixate the tissue. After 30 minutes, the brains were washed 3 times in PBS with 0.3% triton X-100 (PBST) and blocked for 1 hour in 10% goat serum PBST solution with rotation at room temperature. An anti-SNAP25 mouse monoclonal purified IgG antibody (Synaptic Systems, 111 011) was diluted to 1:100 from a 1mg/mL stock solution in the blocking buffer as the primary antibody and added to the blocked brains for overnight incubation at room temperature with rotation. The following day, brains were washed 3 times with PBST and then incubated overnight with an Alexa-647 goat anti-mouse (A11032) secondary antibody diluted 1:1000 in PBST at room temperature. Brains were then washed 3 times in PBST on the final day and mounted for confocal imaging.

### Confocal imaging and image processing

All confocal imaging was performed on a W1 Yokogawa spinning disk attached to a Zeiss Axio Observer Z1 controlled by Slidebook 6.0 software. For co-localisation imaging of SynaptopHluorin and CsChrimson-mCherry, a 20x 0.8 NA Plan-Apochromat and 40x 1.2 NA C-Apochromat water immersion objective was used to image neural structures. For co-localisation imaging of mVenus and SNAP25, the same 20x and a 63x 1.4 NA Plan-Apochromat oil immersion objective was used. All z-slice steps were set to 0.1µm with a variable exposure time of 250 – 1000msec depending on the fluorophore and sample. mVenus and SynaptopHluorin were excited with a 488nm laser wavelength and Alexafluor-647 bound to anti-SNAP25 was excited with a 640nm laser wavelength. All images were captured on a Hamamatsu ORCA-Flash 4.0 V2 sCMOS camera.

For 20x and 40x water imaging experiments, both z-stacks had an average z-projection generated and colour channels merged using ImageJ. For the 63x SNAP25 and mVenus co-localisation experiments, each z-slice in the mVenus stack was subjected to a background subtraction with a 5 pixel (0.5 micron) rolling radius and an ‘IsoData’ auto-threshold (Calvard and Ridler, 1978) to generate a mask for neuronal structure (Figure S6). This mask was converted into an ROI selection, saved, and then used to ‘Clear Outside’ of the ROI. The ROI selection from mVenus expression was registered to the equivalent z-slices in the SNAP25 imaging to perform the same ‘Clear Outside’ process. From this pre-processing, the z-projection of these stacks would only be in the region where mVenus expression was detected. This was to avoid projecting SNAP25 signal in z-planes outside of the mVenus expression, which would confound co-localisation calculations. After this pre-processing, both the mVenus and SNAP25 stacks were z-projected using the average intensity, colour merged, and had their co-localisation R^2^ calculated with the ‘Colocalization threshold’ plugin in ImageJ (Figure S6).

### Data and statistical analysis

Data is presented as the mean ± the standard error of the mean. Individual data points indicate single recordings from multiple brains. For all SynaptopHluorin and sptPALM experiments, recordings were taken on both the left- and right-hand side of the brain due to the bilateral expression patterns in the split-Gal4 lines. Statistical analysis tests for significance were t-tests, one-way ANOVA, and two-way ANOVA (indicated in figure legends where relevant) and performed in Graphpad Prism 9. A significance threshold of 0.05 was employed for all statistical analysis.

## Figure Legends

**Figure S1 –.**
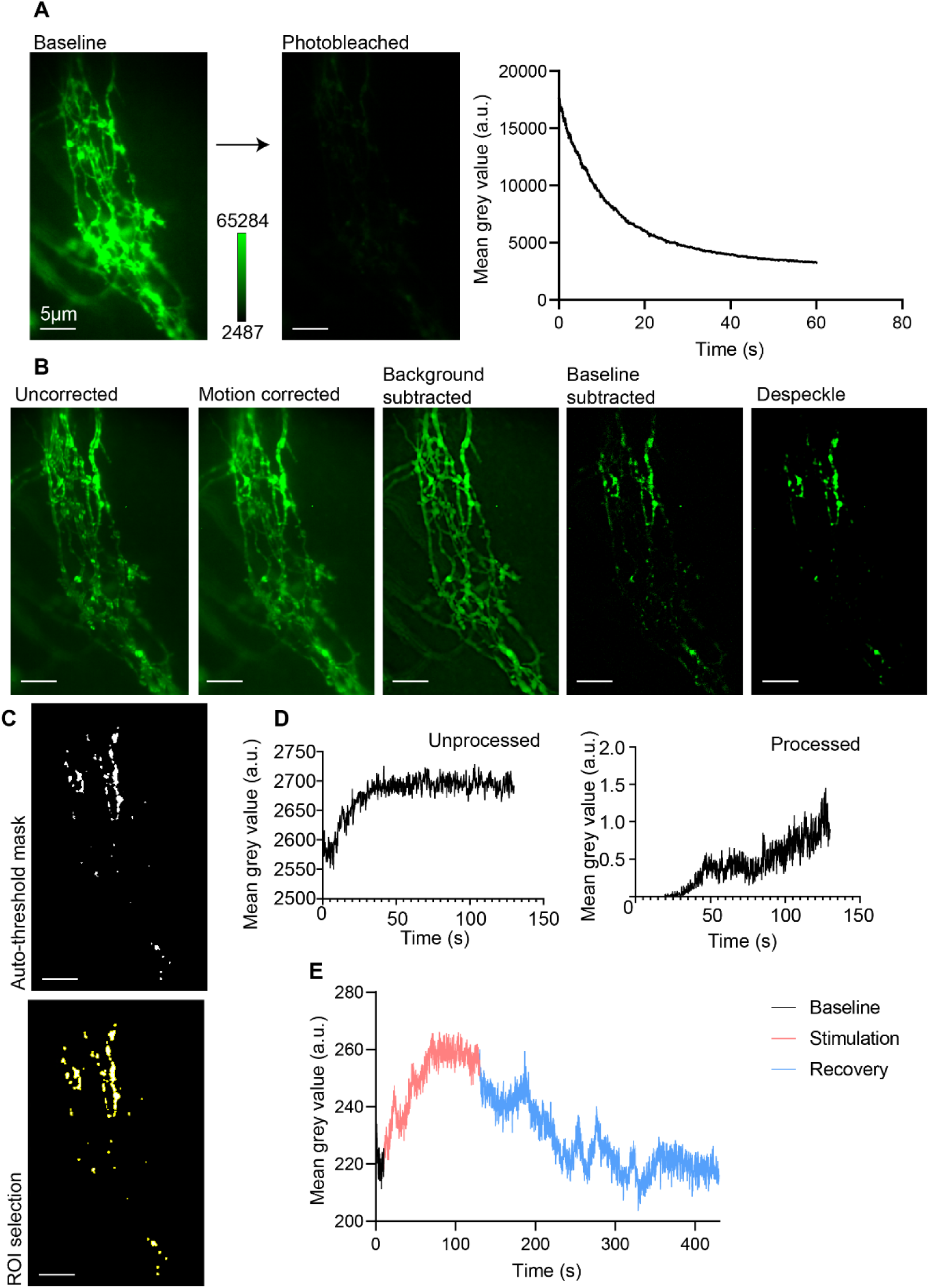
Processing of synaptopHluorin recordings in fly brains. **(A)** SynaptopHluorin expressed under UAS control in cholinergic MB543B-Gal4 neurons. Baseline fluorescence needed to be photobleached prior to experiments due to high background. **(B)** Processing the baseline and stimulation segments of the synaptopHluorin imaging required multiple steps to resolve true changes in fluorescence. Recordings were drift corrected for only the baseline and stimulation due to inaccurate correction with the addition of recovery. Drift corrected movies were background subtracted, baseline subtracted, and then despeckled to eliminate any noise or baseline fluorescence. Images are standard deviation z-projections. **(C)** A ‘Li’ autothreshold algorithm was applied to isolate the highest delta fluorescence changes from a standard deviation z-projection of the final processed recording and ROIs were converted into an ROI selection. **(D)** Comparison between the fluorescence of unprocessed vs processed recordings for baseline and stimulation. **(E)** Measurement of the recovery period of the recording without drift correction and less specific ROI selection shows that synaptopHluorin fluorescence returns to baseline after the red light stimulation is turned off.

**Figure S2.**
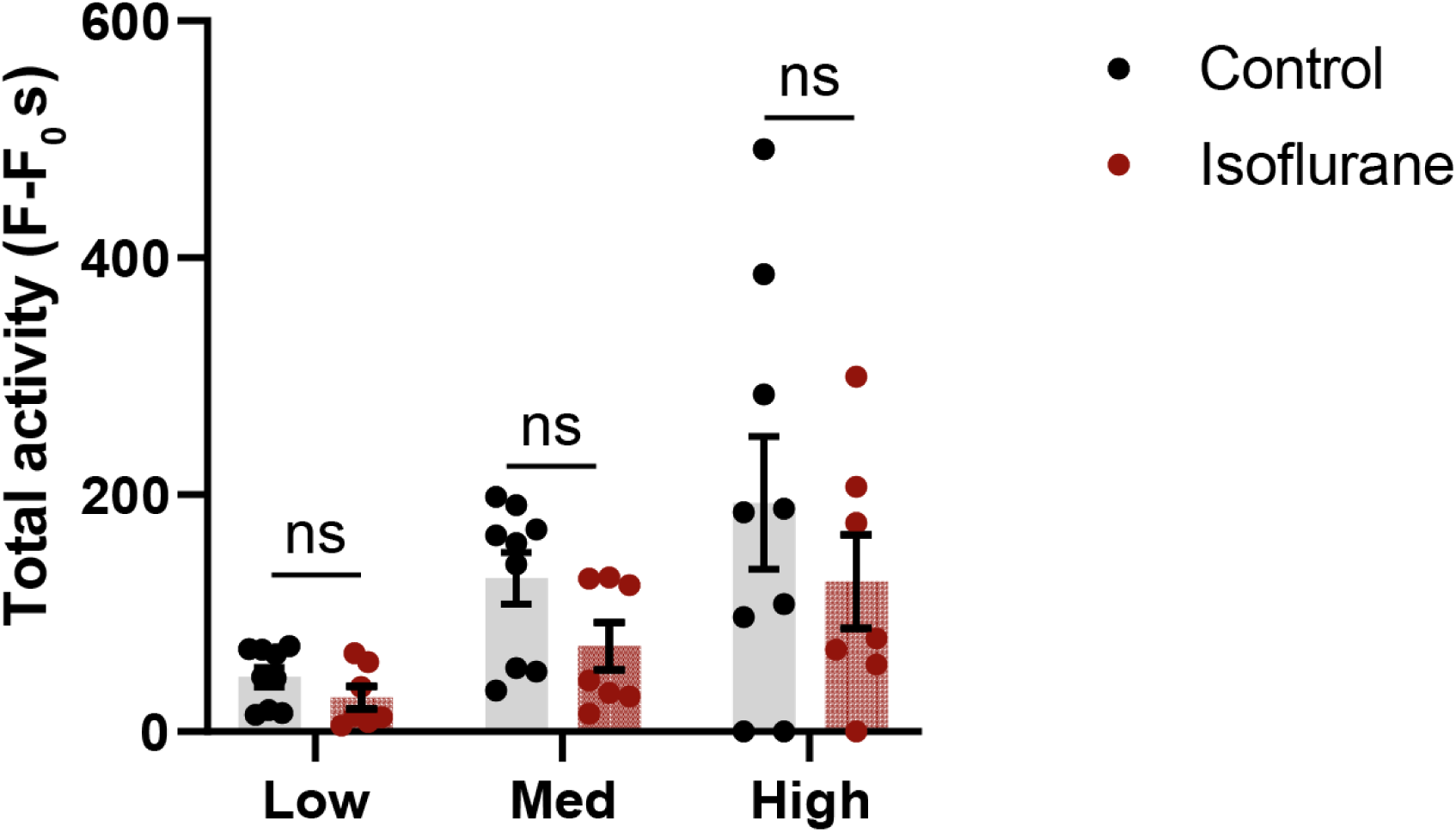
– Total activity of synaptopHluorin unchanged by isoflurane. Total activity of synaptopHluorin (not normalised to the total release area) shows no significant difference between control and isoflurane between the small, medium, and large release sites

**Figure S3 –.**
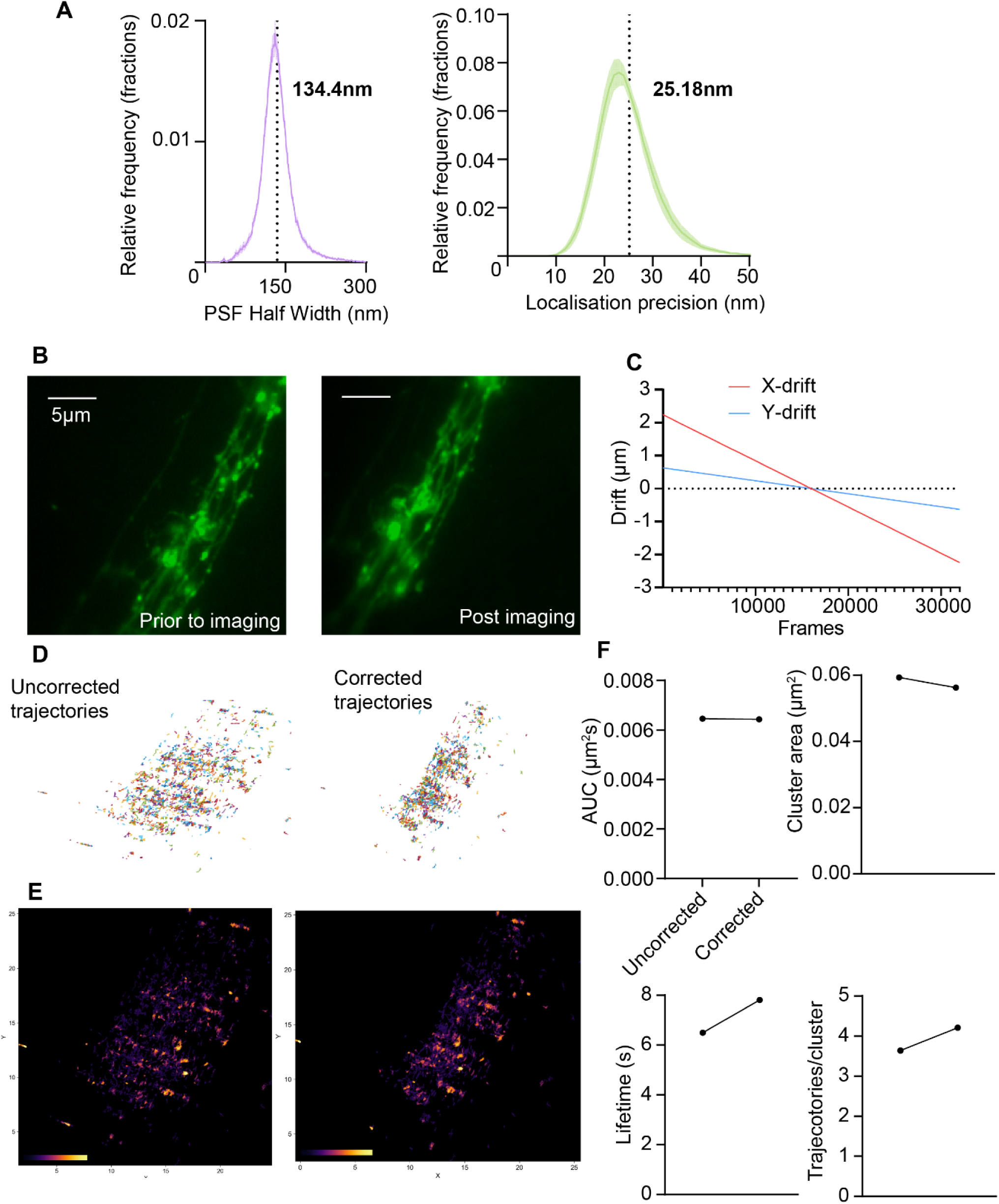
Drift correction of mEos3.2 trajectories for spatiotemporal clustering. **(A)** mEos3.2 molecules were localised with a point-spread function half width of 134.4nm and a localisation precision of 25.18nm. **(B)** Low resolution mEos3.2 images of MB543B-Gal4 cholinergic synapses prior (left) and post (right) imaging shows drift that is incurred. **(C)** Measurements of x- and y-drift, values used to register against trajectory frames and correct drift of molecules post-hoc. **(D)** Trajectories uncorrected (left) and corrected (right). **(E)** Trajectory overlap for uncorrected (left) and correct (right), more overlap and brighter spots for the corrected trajectories. **(F)** Mobility data (MSD, AUC) unaltered by drift correcting trajectories, however differences observed for spatiotemporal clustering metrics.

**Figure S4 –.**
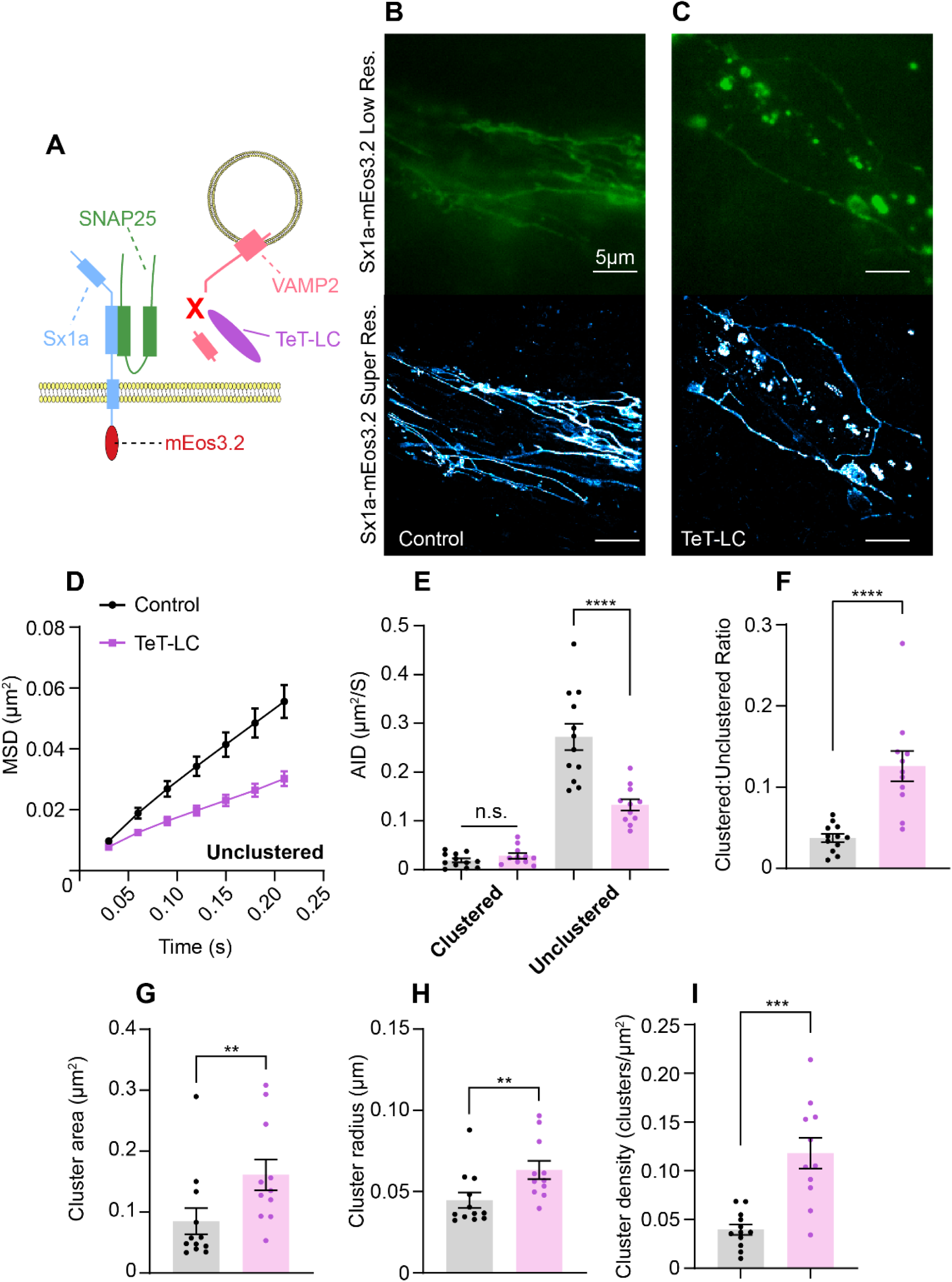
TeT-LC disruption of cholinergic synapse development impairs Sx1a-mEos3.2 mobility. **(A)** Schematic of the SNARE complex with syntaxin1A-mEos3.2 (Sx1a-mEos3.2) and TeT-LC cleavage of VAMP2. **(B)** Expression patterns of Sx1a-mEos3.2 within the synapses of the MB543B-Gal4 cholinergic mushroom body output neuron (MBON) in low resolution (top) and super resolved (bottom) in 3-5 day old female fruit fly brains (scale bar 5µm). **(C)** Expression pattern of Sx1a-mEos3.2 as in B but with developmental TeT-LC expression (scale bar 5µm). **(D)** Mean squared displacement (MSD) analysis of Sx1a-mEos3.2 for unclustered trajectories reveals a significant decrease in mobility with TeT-LC cleavage of VAMP2, resulting from disordered synaptic organisation (Control 0.055 ± 0.005 µm2, n = 12 TeT-LC 0.030 ± 0.002 µm2, n = 11). **(E)** The average instantaneous diffusion (AID) of both clustered and unclustered Sx1a-mEos3.2 trajectories reveals a significant decrease in diffusion of the unclustered population without affecting the mobility within clusters (Clustered control 0.019 ± 0.004 µm2/s TeT-LC 0.028 ± 0.006 µm2/s, n.s. p = 0.2351. Unclustered control 0.272 ± 0.027 µm2/s TeT-LC 0.133 ± 0.012 µm2/s, **** p < 0.0001). **(F)** The number of clustered to unclustered trajectories significantly increased with TeT-LC (Control 0.037 ± 0.005 TeT-LC - 0.126 ± 0.019, **** p < 0.0001). **(G, H, & I)** Alongside an increase in cluster number, TeT-LC also significantly increased the cluster area, radius, and density of Sx1a-mEos3.2 of trajectories in cholinergic synapses (** p < 0.01 *** p < 0.001).

**Figure S5 –.**
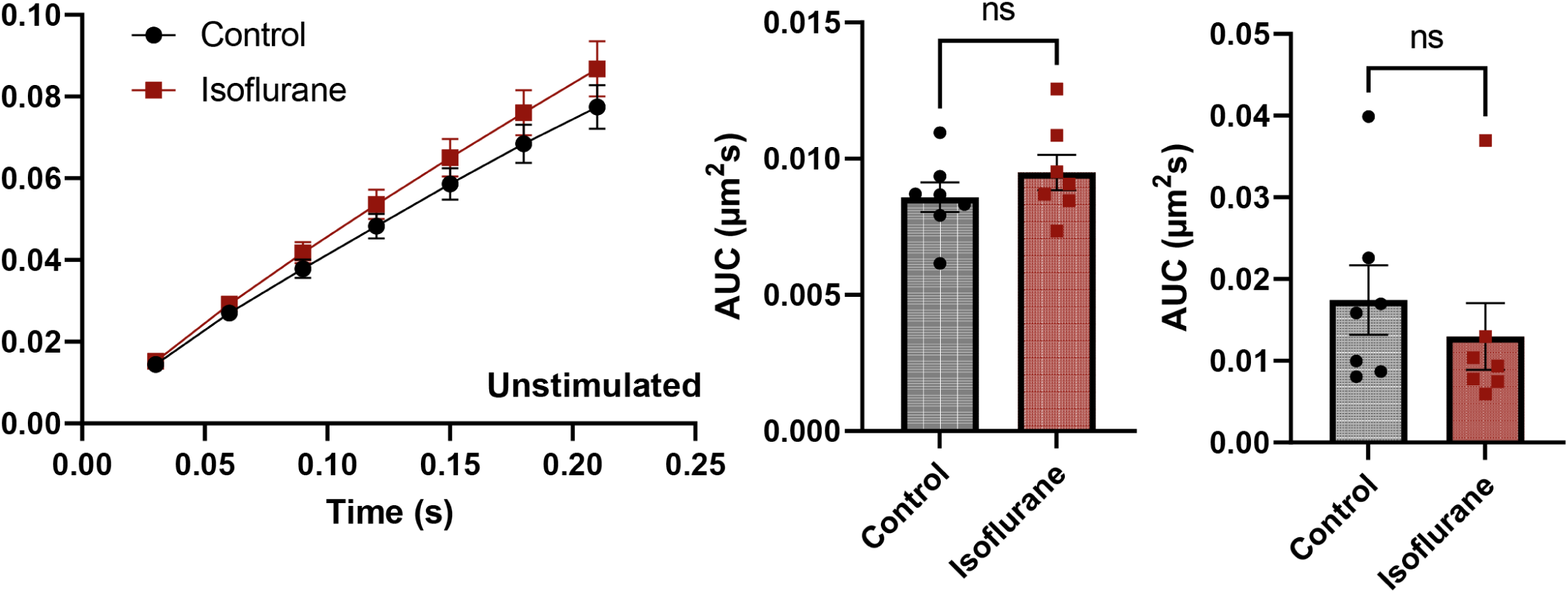
Sx1a-mEos3.2 dynamics unaffected by isoflurane in unstimulated synapses. Flies only expressing UAS-Sx1a-mEos3.2 (without UAS-CsChrimson-mVenus expression). In the presence of isoflurane, Sx1a-mEos3.2 diffusion was unable to be restricted and the clustering unaffected (n = 7 control and isoflurane, Mann-Whitney test).

**Figure S6 –.**
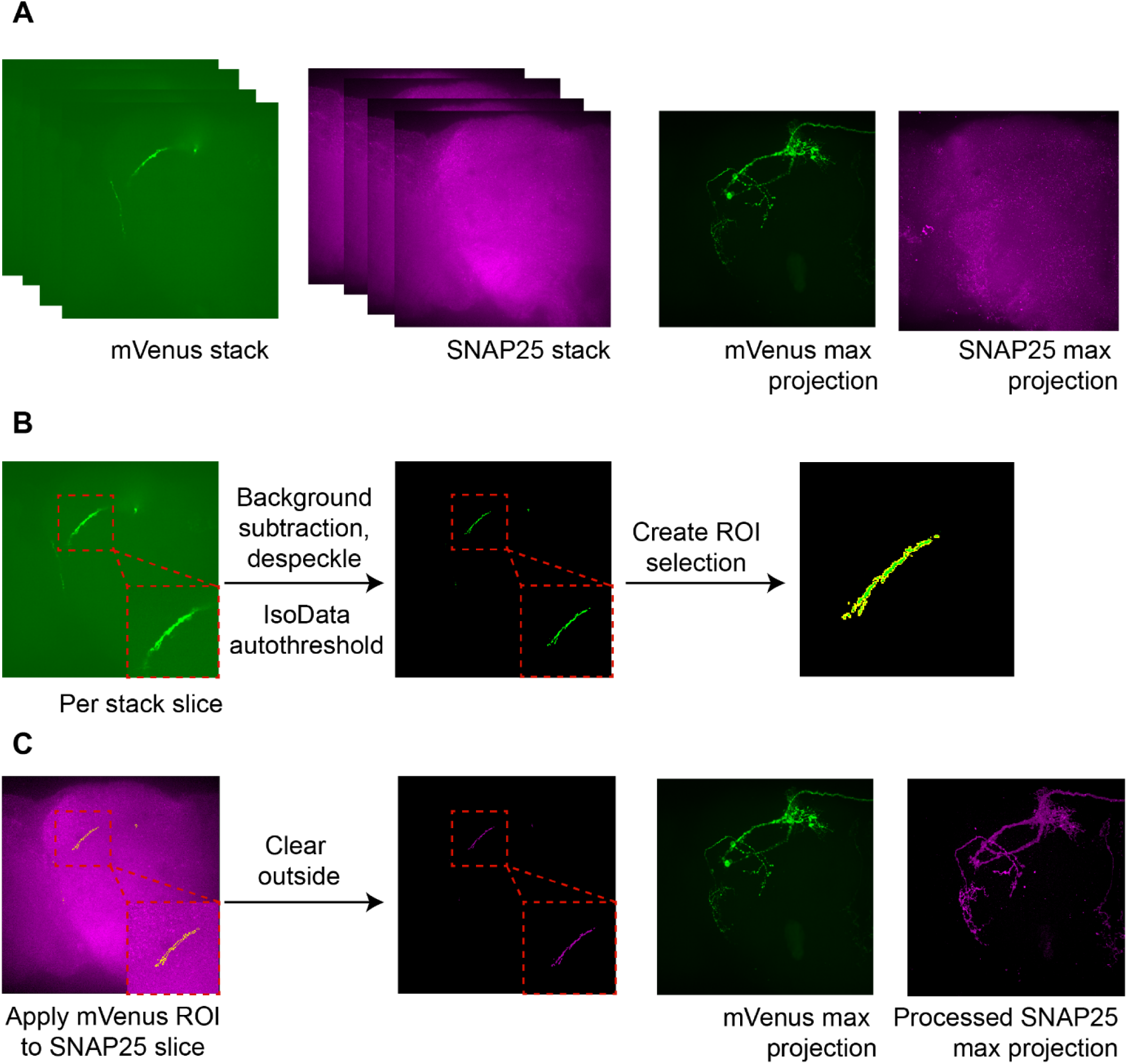
Processing confocal stacks to isolate SNAP25 signal in multiple z-planes. **(A)** mVenus and SNAP25 stacks with maximum z-projection shows that the mVenus signal shows clear expression in cholinergic MB543B-Gal4 compartments, however SNAP25 immunohistochemistry labels abundantly across multiple z-planes **(B)** Each slice of the mVenus stack was background subtracted, despeckled, and had an ‘IsoData’ autothreshold applied to isolate neuronal segments that fluoresce highly. This autothreshold was converted into an ROI selection and stored for every slice in the stack. **(C)** Each slice of the SNAP25 stack had an ROI applied to it from the corresponding mVenus slice with any signal outside of the ROI cleared. The final SNAP25 z-projection stack is signal in an area that only corresponds to detected mVenus signal for each z-plane.

